# Frequency matters: Up- and Down-Regulation of Dopamine Tone Induces Similar Frequency Shifts in Cortico-Basal Ganglia Beta Oscillations

**DOI:** 10.1101/2020.12.14.422621

**Authors:** L. Iskhakova, P. Rappel, G. Fonar, O. Marmor, R. Paz, Z. Israel, R. Eitan, H. Bergman

## Abstract

Beta oscillatory activity (13-30Hz) is pervasive within the cortico-basal ganglia (CBG) network. Studies in Parkinson’s disease (PD) patients and animal models suggested that beta-power increases with dopamine depletion. However, the exact relationship between oscillatory power, frequency and dopamine-tone remains unclear. We recorded neural activity in the CBG network of non-human-primates (NHP) while acutely up- and down-modulating dopamine levels. Further, we assessed changes in beta oscillations of PD patients following acute and chronic changes in dopamine-tone. Beta oscillation frequency was strongly coupled with dopamine-tone in both NHPs and human patients. In contrast, power, coherence between single-units and LFP, and spike-LFP phase-locking were not systematically regulated by dopamine levels. These results demonstrate via causal manipulations that frequency, rather than other properties, is the key property of pathological oscillations in the CBG networks. These insights can lead to improvements in understanding of CBG physiology, PD progression tracking and patient care.

## Introduction

Oscillatory behavior in different frequency bands is common in the cortico-subcortical circuits that loop through the basal ganglia (BG). Activity in the beta range (13-30Hz) has been suggested to play a role in a number of cognitive and motor behaviors. Beta activity is commonly thought to encode the promotion of a “status quo” by maintaining an active process that preserves the current motor directive at the cost of switching to a new one^1^. However, the mechanisms that determine and regulate properties of beta oscillatory activity, like frequency, power and coherence, are still not well defined. There is a significant upregulation of the power (amplitude) of beta activity in the BG of Parkinson’s disease (PD) patients^2–4^ and PD animal models^5–7^. In line with the “status quo” hypothesis, PD beta activity has been correlated with akinetic symptoms^8^. Thus, the power of beta activity in the CBG network has been considered a marker for parkinsonism and consequently, many researchers, including our group, focused on studying beta activity amplitude in parkinsonian patients and animal models^5,9^.

The neuropathological hallmark of PD is progressive degeneration of midbrain dopaminergic neurons and their projections to CBG networks. Therefore, a high prevalence of beta oscillations in PD patients and animal models hinted to a potential role for dopamine in generation of beta oscillations. Acute dopamine modulation in rodent animal models has not always resulted in changes in beta properties^10,11^. However, in 6-OHDA rodents an upregulation of dopamine with apomorphine, a dopamine agonist, resulted in an increase in beta-frequency and a decrease in beta-power^12^. Analysis of PD patients^13^ and MPTP treated NHPs^5,14^ revealed a high power beta activity compared to dopamine treated patients or primate normal controls. In the PD circuitry, the high power of beta activity was accompanied by an increase in coherence within and between CBG networks^6,10,12,15,16^. Both were alleviated by dopamine replacement therapy (DRT) or deep brain stimulation (DBS)^10,12,17–19^. Based on these studies of beta activity in PD patients and animal models, dopamine has been suggested to play an important role in modulating the power of beta signaling in the CBG network. However, because most of this evidence was derived from dopamine-depleted PD patients and animals models we cannot reliably assume that what we gleaned from these studies provides us with a comprehensive depiction of interactions between dopamine tone and beta oscillation properties, including power, frequency, coherence and single-unit entrainment.

To examine whether dopamine tone is regulating beta activity we recorded single-unit activity (SUA) and local field potential (LFP) in the CBG circuit of healthy, awake behaving non-human primates (NHP) after acute dopamine up- and down-modulation using apomorphine (dopamine agonist), amphetamine (dopamine transporter (DAT) inhibitor), and haloperidol (dopamine antagonist). To further examine the effects of acute and chronic dopamine modulation in human subjects we exploited the data collected from PD patients before and after dopamine therapy up to 250 days post-surgery using Activa PC+S (Medtronic Inc., Minneapolis, MN, USA). This allowed us to measure changes in beta properties as a function of acute dopamine modulation and progressive dopamine loss that is a hallmark of PD. With a general goal to assemble a comprehensive and nuanced understanding of beta physiology in the CBG circuitry, our study revealed that beta-frequency, rather than power or any other property, is the key marker of dopamine tone and pathological oscillations in the CBG networks. We further propose that progressive beta-frequency decline, and not a beta-power amplitude, could be used as a more effective indicator of PD progression and as a trigger for adaptive DBS procedures.

## Results

We examined SUA and LFP recordings collected with multiple micro-electrodes from the dorso-lateral prefrontal cortex (dlPFC) and the external segment of the globus pallidus (GPe), the central nucleus of the BG circuitry^5^, of awake, behaving monkeys under acute up- and down-modulation of dopamine tone (Fig.1A-D, Table S1)^20^. Cortical units were separated into putative-pyramidal cells (wide) and putative-interneurons (narrow) based on the width of the spike shape (Fig.S1)^21,22^. Our assessment of human electrophysiology utilized multiple post-surgery recording of bilateral subthalamic nucleus (STN) LFP from 4 PD patients on and off DRT over a period of 170 to 250 days (Fig.1E-H, Table S2).

**Figure 1:**
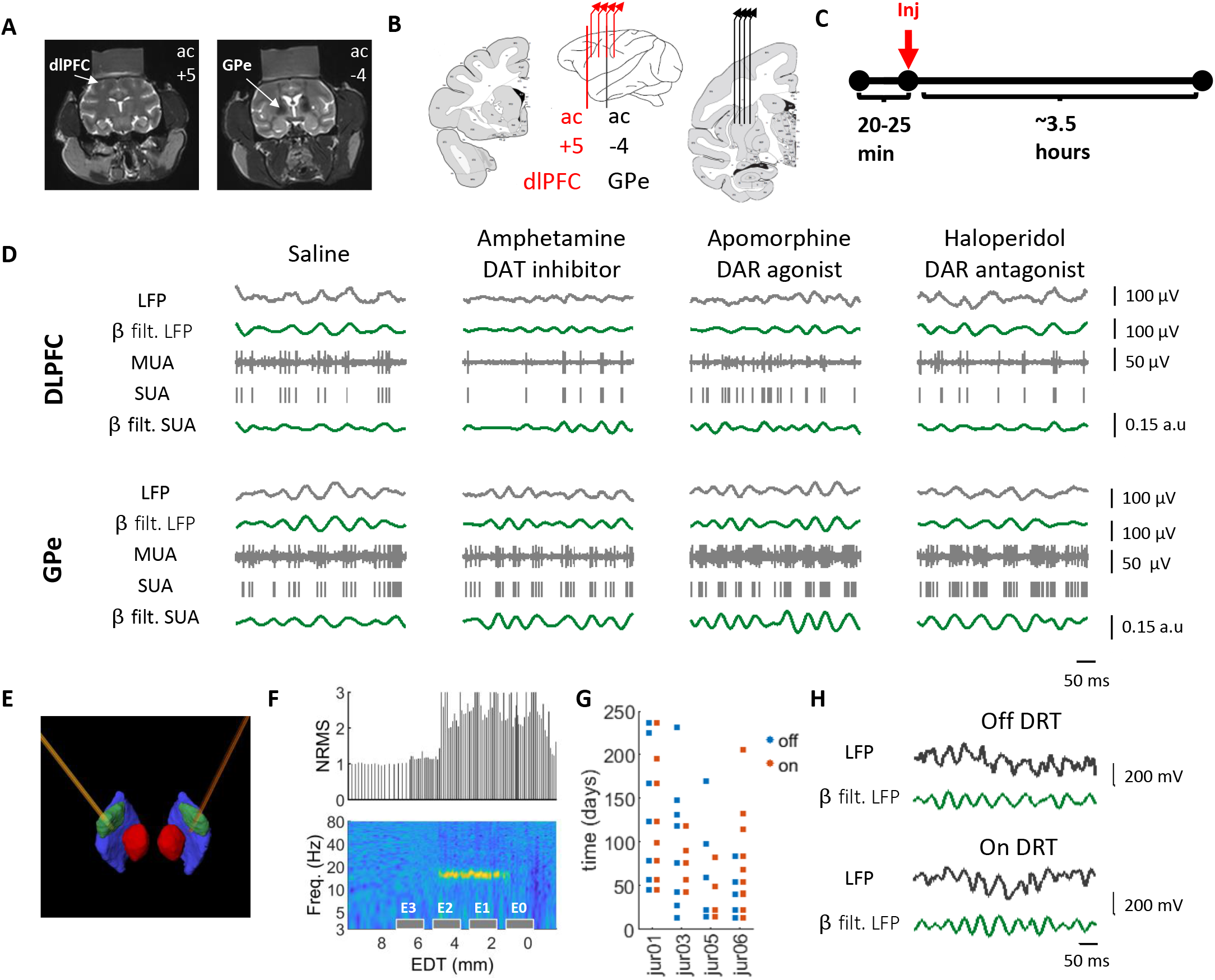
Experiment design. (A) MRI of monkey G. Coronal images showing recording targets. (B) A scheme of ac+5 and ac-4 coronal planes with four electrodes in each recording target. Middle: coronal plane positions marked on an atlas scheme. Adapted from Martin and Bowden^20^. (C) Daily timeline scheme with pre- and post-injection times. (D) 500 ms traces from the dlPFC and GPe under each drug condition. LFP: local field potential, MUA: multiunit activity, SUA: single unit activity (cortical narrow units and pallidal units), β filt: beta (8-24Hz) bandpass filtered. (E) Electrode position marked on reconstruction of one patient atlas, based on the postop CT with the pre-op MRI. Green: STN, red: red nucleus, blue: substantia nigra pars reticulata. (F) Electrode contact position relative to STN electrophysiological activity of the same patient as in (E). x axis indicates estimated distance from clinical target (EDT). The target was set preoperatively to the estimated ventro-lateral border of the STN. Top: MUA total power evaluated as normalized root mean square (NRMS). NRMS elevation and decline indicate STN entry and exit, respectively. Bottom: LFP normalized power spectral density (nPSD, percentage of total power, filtered with gaussian window for presentation purposes). Contact positions marked as grey boxes. The STN dorsal motor area can be identified according to its pronounced beta activity. (G) Recording schedule of PD patients included in this study. (H) 500ms traces from the STN of PD patient (same patient as in E-F) on and off DRT. DAT – dopamine transporter, DAR – dopamine receptor.

### Up- and down-modulation of dopamine tone up- and down-shifts LFP beta-frequency, respectively

The population average spectrogram of LFP signal recorded in the dlPFC and GPe revealed a clear shift in frequency and amplitude of beta oscillations in response to both up- and down-modulation of dopamine tone (Fig.2A). Beta signal from the GPe exhibited greater power in comparison to that from the dlPFC. Saline injections resulted in maintenance of beta oscillation frequency, while acute upregulation of dopamine tone by amphetamine (DAT blocker) shifted the frequency of beta oscillations up and downregulation of dopamine tone by haloperidol (dopamine receptor antagonist) shifted beta-frequency down. Beta shift was maintained for the duration of the recording (~3hours). Apomorphine (dopamine receptor agonist) injection resulted in two distinct phases. At first, similar to amphetamine, frequency of beta oscillations shifted up (Apo1), but about 1 hour after injection beta-frequency depressed to about baseline levels (Apo2). Diverging from the post-amphetamine effect, beta-power decreased during Apo1, and as the beta-frequency declined in Apo2, beta-power was reestablished.

**Fig 2.**
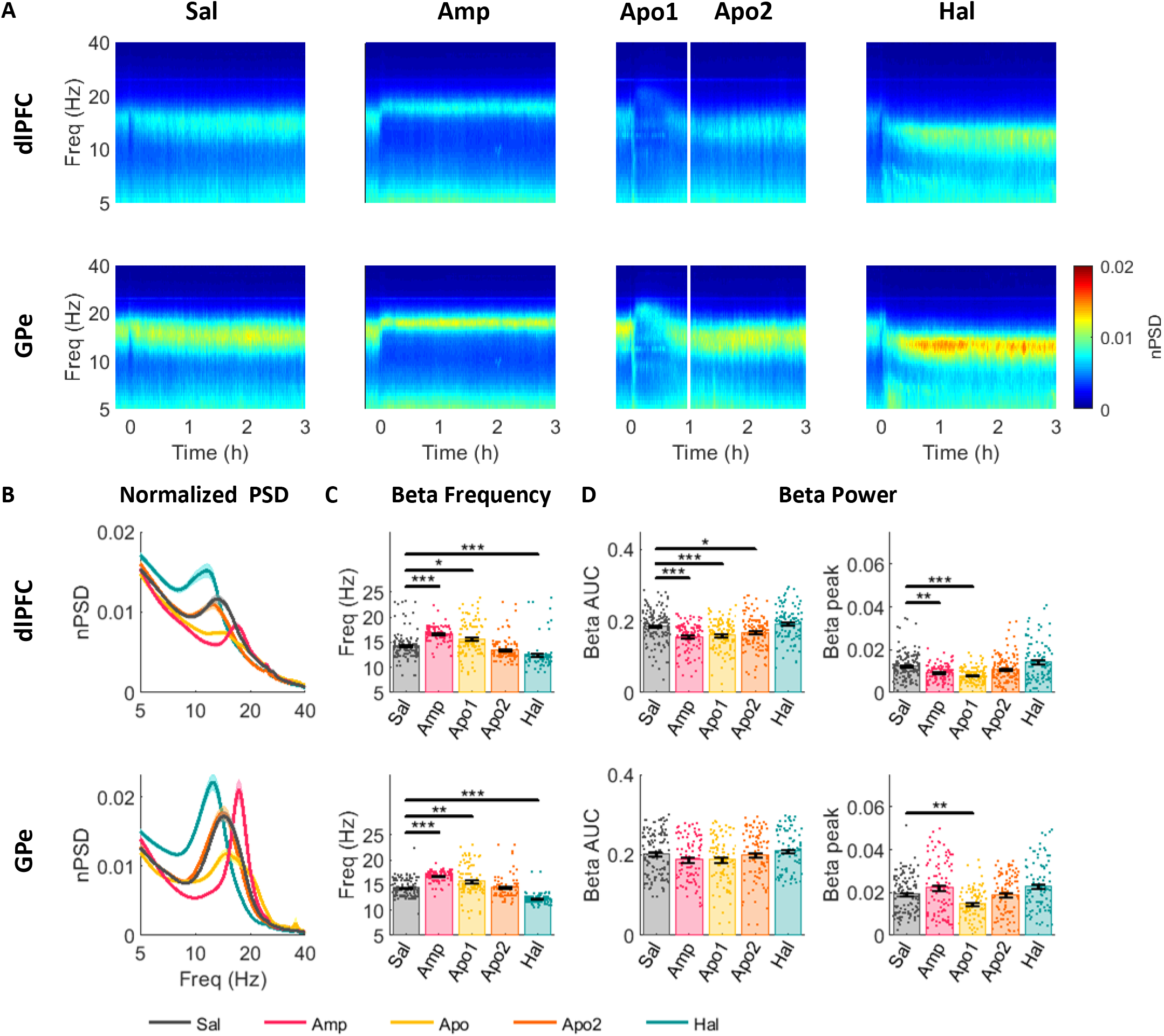
Acute up- and down-modulation of dopamine tone up- and down-shifts LFP beta frequency in the dlPFC and GPe of NHP. (A) Average spectrogram of dlPFC (top) and GPe (bottom) LFP. Time 0 on the x-axis indicates injection time. Color scale indicates nPSD. White line in the third column divides the post-apomorphine period into Apo1 - agonistic phase and Apo2 – post-agonistic phase. (B-D) LFP beta properties in dlPFC (top) and GPe (bottom) under each drug condition. (B) Average nPSD. (C) Frequency of beta peaks in oscillatory LFP sites (see methods). (D) Beta power evaluated as area under curve (AUC) of the nPSD in 8-24Hz range (left), or as nPSD beta peak within 8-24 Hz frequency band (right). Bars indicate average values. Single points indicate individual LFP site values. Black vertical lines indicate standard error of the mean. Drug influence was evaluated by Kruskal-Wallis test followed by post-hoc pairwise comparisons. Comparisons between saline and drug treatments are presented in current figure. Full post-hoc results can be found in Table S3. * p<0.05 ** p<0.01 *** p<0.001

Quantitative analysis of LFP beta oscillation properties (Fig.2B-D, post-hoc results are in Table S3) confirmed that the dopamine tone can bidirectionally shift the frequency of LFP beta oscillations in both cortex (Fig.2B-C; χ^2^(4)=185.13, p=5.9e-39, η^2^=0.23) and GPe (Fig.2B-C; χ^2^(4)=228.69, p=2.5e-48, η^2^=0.35). Control (saline) beta-frequency at about 14Hz in both brain areas was flanked on the left/lower frequency band by the haloperidol injection peak (~12Hz) and on the right/higher frequency by amphetamine and apomorphine (Apo1) injection peaks (~17Hz). Downward shift of beta-frequency in Apo2 recovered the properties of control beta.

Effect of dopamine tone modulation on beta-power was less robust and inconsistent (Fig.2D). Beta-power was measured by two methods, first by the area under curve (AUC) which estimated the total power within the beta band (8-24Hz) and second, by the amplitude of the peak. In the dlPFC, up-modulation of dopamine tone by amphetamine or apomorphine (Apo1) resulted in decreased beta-power relative to control (beta-AUC: χ^2^_(4)_=55.54, p=2.5e-11, η^2^=0.09; beta-peak: χ^2^_(4)_=59.36, p=3.9e-12, η^2^=0.12). Haloperidol-induced up-modulation of beta-power was not significantly different from control in the dlPFC. The effect of dopamine tone up-modulation on beta-power was not consistent in the GPe (beta-AUC: χ^2^_(4)_=8.98, p=0.06, η^2^=0.02; beta-peak: χ^2^_(4)_=30.02, p=4.8e-6, η^2^=0.07). Apomorphine (Apo1), but not amphetamine, decreased beta-power relative to control. Analysis of the highest 20% of beta-peak values in each drug category showed a significant effect of drug treatment on beta-power within this subsection (χ^2^_(4)_=72.85, p=5.7e-15, η^2^=0.69). In comparison with saline, beta-power was higher post-amphetamine (p=1.6e-4, hedge’s g=−1.92) and haloperidol (p=0.002, hedge’s g=−1.45) and lower post-apomorphine (Apo1) (p=0.03, hedge’s g=−1.73), suggesting that in some cases both up- and down-modulation of dopamine tone can induce increases in beta-power.

### Up- and down-modulation of dopamine up- and down-shifts beta-frequency of spiking activity, respectively

Initial test of single-unit firing rate (FR) confirmed that, as previously reported^9^, dopamine up-and down-modulation alters the FR of single-units in the CBG (Fig.S2; wide units: χ^2^(4)=31.56, p=2.4e-6, η^2^=0.008; pallidal units: χ^2^(4)=80.94, p=1.1e-16, η^2^=0.05). Amphetamine increased the FR of cortical wide units relative to control (p=2.8e-4, hedge’s g=−0.14). FR of cortical narrow units was not significantly modulated by dopamine tone (χ^2^(4)=7.78, p=0.1, η^2^=0.02). FR in GPe units increased after amphetamine (p=0.023, hedge’s g=−0.22) and apomorphine (Apo1; p=9.4e-7, hedge’s g=−0.55) injection and decreased after haloperidol injection relative to control (p=6.0e-4, hedge’s g=0.29). Once the single-unit FRs were confirmed to follow previously established patterns, the SUA was examined for expression of beta oscillations.

In agreement with LFP results, acute dopamine tone modulation up/down-shifted the frequency of beta oscillations in narrow cortical units (χ^2^(4)=18.35, p=0.001, η^2^=0.06) and pallidal units (χ^2^(4)=52.36, p=1.2e-10, η^2^=0.02), but not in the wide cortical units (χ^2^(4)=2.35, p=0.67, η^2^=0.001) (Fig.3A,B, post-hoc results are in Table S4). Haloperidol-induced low dopamine tone resulted in a lower frequency of beta oscillations in comparison to control in both narrow cortical and pallidal units. An increase in dopamine tone post-amphetamine shifted the frequency of pallidal beta oscillations to a higher frequency range.

**Fig 3.**
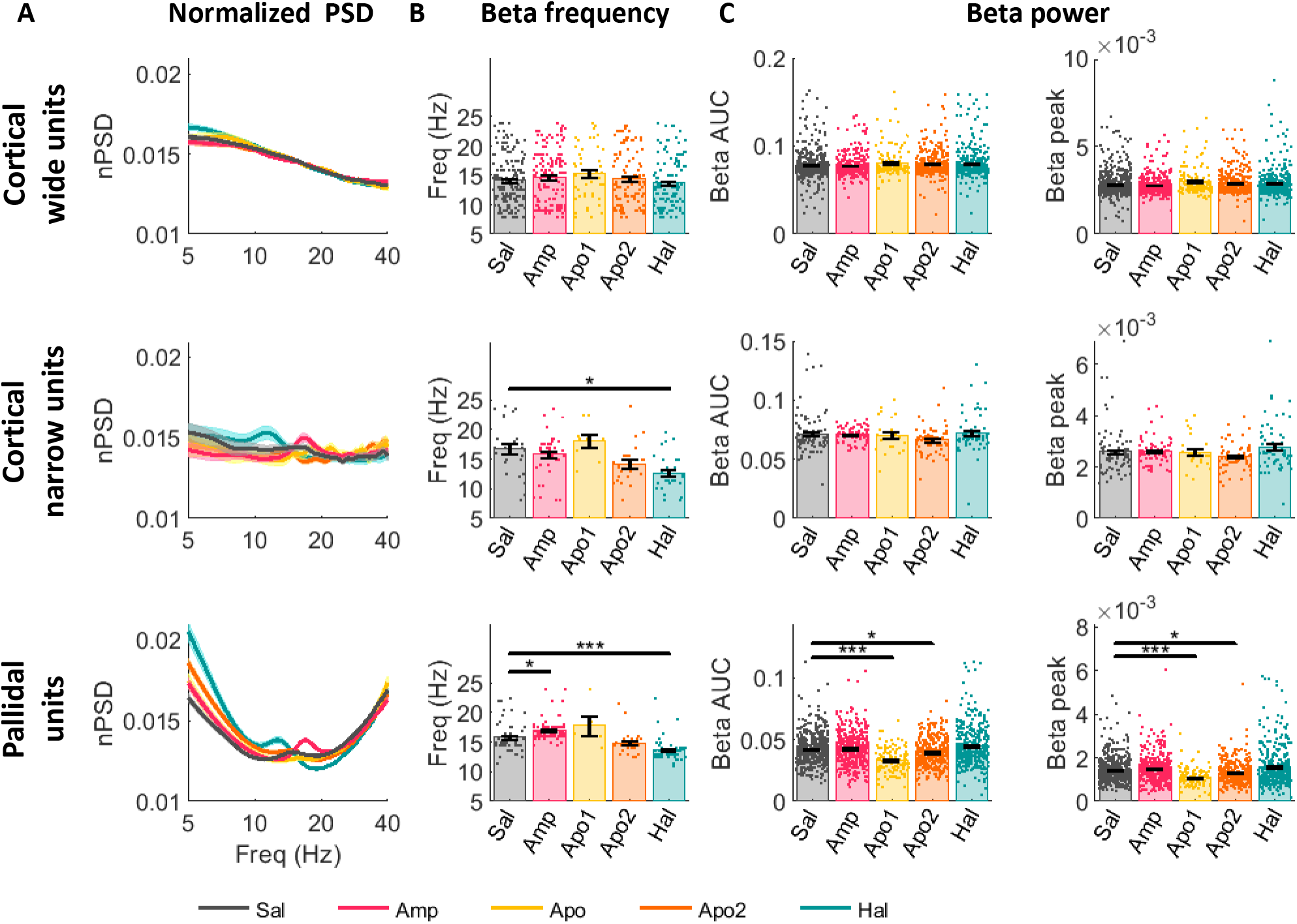
Acute up- and down-modulation of dopamine tone up- and down-shifts SUA beta frequency in the cortical narrow and pallidal units of NHP. Single unit beta properties in cortical wide (top), narrow (middle) and pallidal (bottom) units under each drug condition. (A) Average nPSD. (B) Frequency of beta peaks in oscillatory units (see methods). (C) Beta power evaluated as area under curve (AUC) of the nPSD in 8-24 Hz range (left), or as nPSD beta peak within 8-24 Hz frequency band. Bars indicate average values. Single points indicate individual unit values. Black vertical lines indicate standard error of the mean. Drug influence was evaluated by Kruskal-Wallis test followed by post-hoc pairwise comparisons. Comparisons between saline and drug treatments are presented in current figure. Full post-hoc results can be found in Table S5. * p<0.05 ** p<0.01 *** p<0.001

The power of beta oscillations in pallidal units was modulated by dopamine tone (beta-AUC: χ^2^(4)=76.72, p=8.6e-16, η^2^=0.05; beta-peak: χ^2^(4)=64.29, p=3.6e-13, η^2^=0.04), but the relationship between dopamine tone and beta-power was not monotonic. Apomorphine (Apo1 and Apo2), but not amphetamine, induced a significant decrease in beta-power compared to control (Fig.3C). Both AUC and beta-peak analyses for haloperidol and amphetamine showed no statistical difference from control (Fig.3A,C). However, amphetamine significantly increased the number of oscillatory cortical narrow (χ^2^(4)=11.70, p=0.0197, Cramér’s V=0.19) and pallidal (χ^2^(4)=38.75, p=7.8e-8, Cramér’s V=0.15) units (Fig.S3, post-hoc at Table S5).

LFP and SUA in saline condition were also compared to that of the naïve animals, to test for an effect of injection and recording process (Fig.S4,5). In both cortical (t_(363)_=6.19, p=1.6e-9, hedge’s g = 0.71) and pallidal (t_(508)_=4.77, p=2.4e-6, hedge’s g = 0.42) LFP, beta oscillations in the saline condition had a lower frequency relative to that of naïve animals. Pallidal units’ beta-power was higher in the saline condition (beta-AUC: t_(1551)_=−3.74, p=2e-4, hedge’s g=0.21; beta-peak: t_(1551)_=−4.13, p=3.8e-05, hedge’s g=0.21). These effects could be explained as an outcome of minimal damage caused by chronic microelectrode recording and/or a mild dopamine depletion throughout the experiment period. However, no frequency shift was seen in single-unit oscillations between naïve and saline conditions.

### Frequency of beta LFP coherence in the dlPFC and GPe is shifted by acute up- and down-modulations of dopamine tone

Coherence is used to measure synchrony within and between neural populations. Up- and down-regulation of dopamine tone shifted the central frequency of LFP coherence in the beta range (Fig.4A-C, post-hoc results are in Table S6) within the dlPFC (χ^2^(4)=281.07, p=1.3e-59, η^2^=0.18), GPe (χ^2^(4)=399.93, p=2.9e-85, η^2^=0.47), and between the dlPFC and the GPe (χ^2^(4)=213.52, p=4.6e-45, η^2^=0.37), in the same direction as for the single site (Fig.2 and 3) beta oscillations. Haloperidol down-shifted the coherence peak frequency, while apomorphine and amphetamine up-shifted the coherence peak frequency within the beta range. Interestingly, while cortical LFP coherence within hemispheres was greater than that between hemispheres, in the GPe, coherence within and between hemispheres was comparable (Fig.S6). On the single-unit level, pallidal-pallidal and dlPFC narrow-narrow unit pairs exhibited dopamine tone dependence in coherence beta-frequency (Fig.S7).

**Fig 4.**
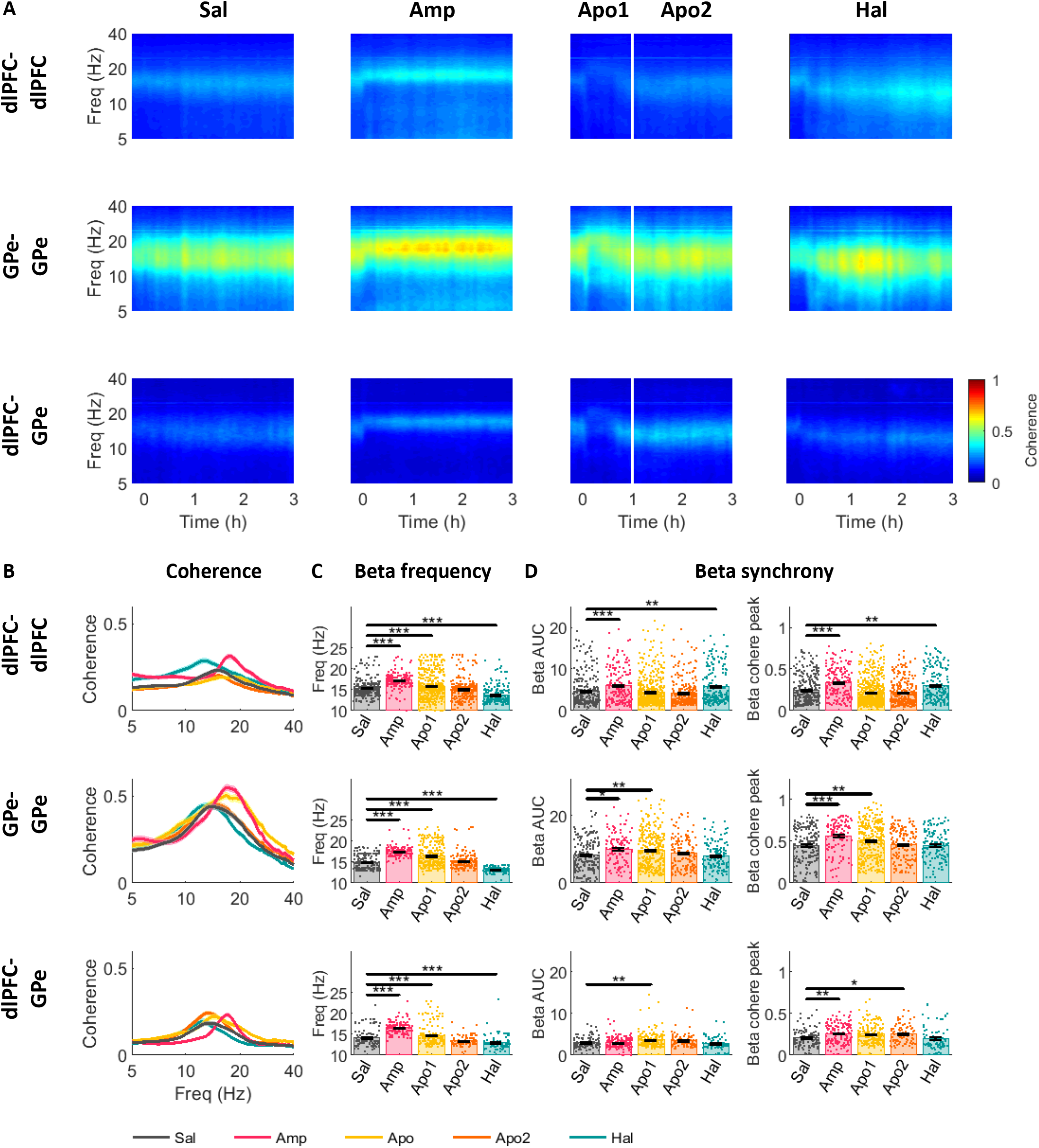
Acute up- and down-modulation of dopamine tone up- and down-shifts LFP beta coherence frequency in the CBG network of NHP. (A) Average coherogram of dlPFC-dlPFC (top), GPe-GPe (middle) and dlPFC-GPe (bottom) LFP pairs. Time 0 on x-axis indicates injection time. Color scale indicates coherence. White line in the third column divides the post-apomorphine period into Apo1 - agonistic phase and Apo2 – post-agonistic phase. (B-D) LFP beta coherence properties in dlPFC-dlPFC (top), GPe-GPe (middle) and dlPFC-GPe (bottom) LFP pairs under each drug condition. (B) Average coherence. (C) Frequency of beta coherence peaks in synchronized LFP sites (see methods). (D) Beta synchrony evaluated as area under the curve (AUC) of coherence in 8-24Hz range (left), and as coherence peak within 8-24Hz frequency band. Bars indicate average values. Single points indicate individual LFP pair values. Black vertical lines indicate standard error of the mean. Drug influence was evaluated by Kruskal-Wallis test followed by post-hoc pairwise comparisons. Comparisons between saline and drug treatments are presented in current figure. Full post-hoc results can be found in Table S6. * p<0.05 ** p<0.01 *** p<0.001

Up- and down-modulation of dopamine tone perturbed the degree of LFP synchrony within the beta range between dlPFC-dlPFC (beta-AUC: χ^2^(4)=43.64, p=7.6e-9, η^2^=0.03; beta-peak: χ^2^(4)=75.84, p=1.3e-15, η^2^=0.06), GPe-GPe (beta-AUC: χ^2^(4)=26.55, p=2.4e-5, η^2^=0.05; beta-peak: χ^2^(4)=41.09, p=2.6e-8, η^2^=0.05), and dlPFC-GPe (beta-AUC: χ^2^(4)=26.63, p=2.4e-5, η^2^=0.05; beta-peak: χ^2^(4)=27.78, p=1.4e-5, η^2^=0.04) sites (Fig.4D, post-hoc results are in Table S6).

In the dlPFC, LFP beta synchrony increased after dopamine up-modulation by amphetamine and down-modulation by haloperidol (Fig.4D). In the GPe, LFP beta coherence was only increased by dopamine up-modulation by amphetamine or apomorphine (Apo1), but not by dopamine down-modulation. LFP beta coherence between dlPFC and GPe was also increased by dopamine up-modulation by amphetamine and apomorphine (Apo1) and in the recovery period after apomorphine injection (Apo2). Importantly, apomorphine first induced a reduction in dlPFC-GPe beta coherence (Fig.4A), similar to its effect on beta-power (Fig.2). The significant enhancement seen during Apo1 (Fig.4D) is probably due to some overlap between Apo1 and Apo2. Analysis of LFP-LFP phase-locking value (PLV) revealed a similar shift in the frequency of beta synchronization (Fig.S8, post-hoc results are in Table S7).

### Acute up- and down-modulation of dopamine tone redirects spike-LFP entrainment to opposing beta phases

Dopamine modulation changed the number of units entrained to LFP beta oscillations in cortical wide (χ^2^(4)=51.87, p=1.5e-10, Cramér’s V=0.18), narrow (χ^2^(4)=10.95, p=0.027, Cramér’s V=0.198) and pallidal (χ^2^(4)=51.52, p=1.7e-10, Cramér's V=0.19) units (Fig.5A; post-hoc results are in table S8). Up-modulation of dopamine tone by amphetamine increased the number of entrained wide cortical units, while apomorphine (Apo2) led to a reduction in entrained units. In the GPe, haloperidol increased the number of entrained units relative to control, while apomorphine (Apo1) decreased it. Results in narrow cortical units showed similar pattern to pallidal units, though comparisons did not yield a significant difference between control and drug conditions.

**Fig 5.**
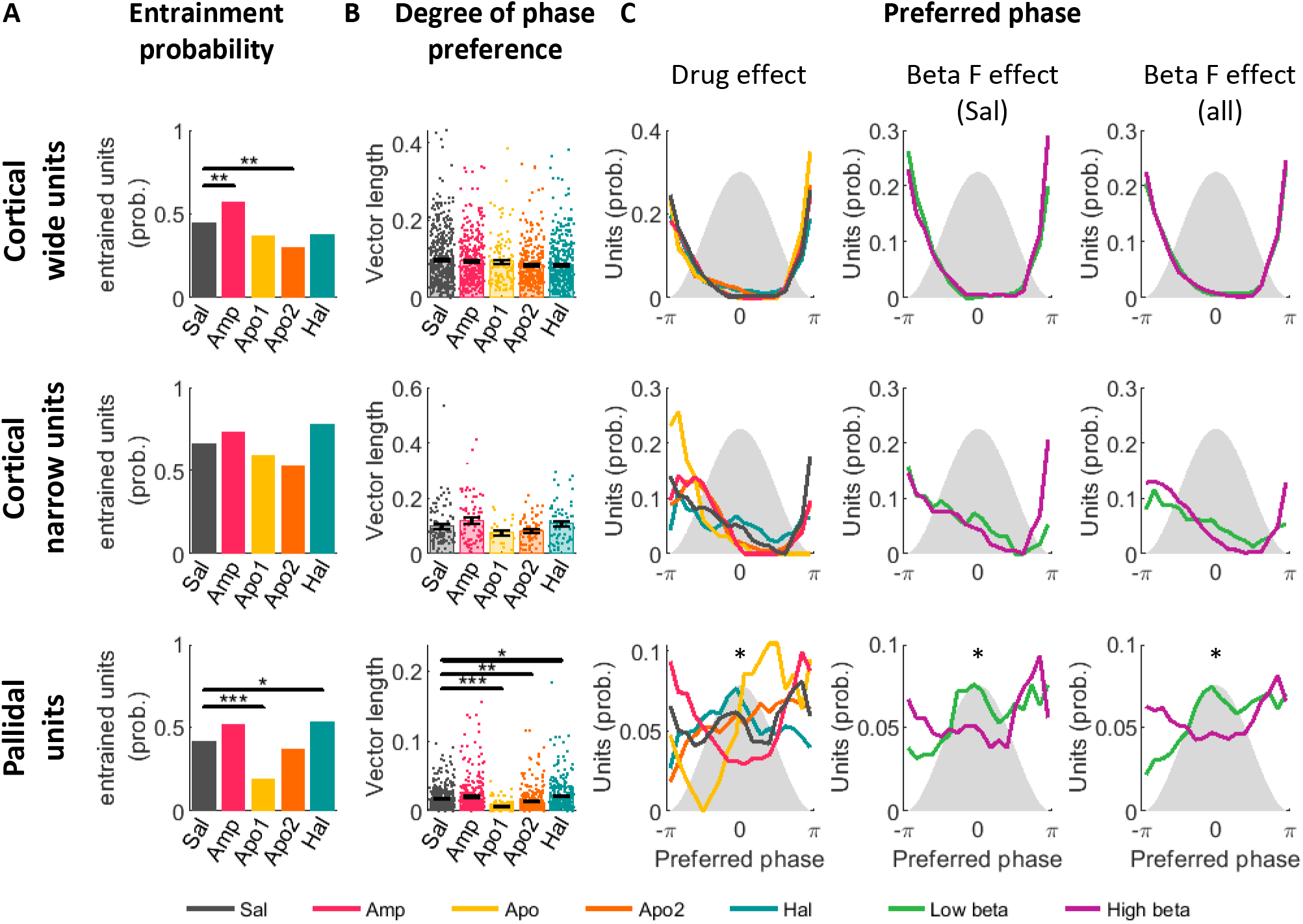
Drug-induced LFP beta frequency affects preferred phase of entrained units. Properties of unit-LFP entrainment in the beta range frequency band of cortical wide (top), narrow (middle) and pallidal (bottom) units under each drug condition. (A) Probability of entrained units. (B) Degree of phase preference was assessed per unit by the vector length of the spike phase circular average. Bars indicate average values. Single points indicate individual unit values. Black vertical lines indicate standard error of the mean. Drug influence was evaluated by Kruskal-Wallis test followed by post-hoc pairwise comparisons. Comparisons between saline and drug treatments are presented in current figure. Full post-hoc results can be found in Table S7. (C) Preferred phase of entrained units. Gray shadow represents LFP beta cycle and x-axis indicates LFP beta phase. Y-axis indicates unit probability to lock to a given phase. Drug influence was evaluated by circular median test followed by post-hoc comparisons when needed. Left: units are grouped by drug condition. Middle: Only saline units, grouped by the unit beta frequency. Right: units from all drug conditions, grouped by the unit beta frequency. In middle and right columns, units were segregated into low-beta and high-beta groups according to the unit beta frequency using a 15Hz cutoff. * p<0.05 ** p<0.01 *** p<0.001

The degree of spike-to-LFP entrainment can be assessed by the vector-length of the spike phase circular average. A large vector-length indicates a high tendency of spikes to cluster around a specific phase of the LFP oscillation. Modulation of dopamine tone affected the vector-length in pallidal units (Fig.5B, post-hoc results are in Table S8; χ^2^(4)=99.83, p=1.1e-20, η^2^=0.04). Down-regulation of dopamine tone by haloperidol increased the vector-length relative to control. Upregulation of dopamine tone by apomorphine (Apo1) decreased the vector-length relative to control. During the post-agonistic period (Apo2) vector-length recovered and exceeded control values. Amphetamine, however, did not generate a similar effect to apomorphine. In cortical wide units our results indicated a significant effect of dopamine modulation on vector-length (Fig.5B; χ^2^(4)=11.02, p=0.026, η^2^=0.009), but post-hoc comparisons failed to reveal a significant effect of drug treatments, although the difference between saline and haloperidol approached significance (Table S8; p=0.054). In cortical narrow units, the trend of results was similar to that seen in pallidal units, but with no statistical significance (Fig.5B).

Next, we focused on the effect of drug injection on the preferred phase of entrained units (Fig.5C left, post-hoc results are in Table S8). These results showed that cortical wide and narrow units maintained their phase-locking to the trough of the beta cycle throughout all the different acute dopamine modulations. In contrast, in pallidal units, circular median test revealed a significant modulation of preferred phase by drugs (P=33.45, p=9.7e-07), but post-hoc comparisons failed to reveal the origin of this effect (Table S8) probably due to low statistical power of the statistical test available to use for this data type. A visual inspection suggested that in the control condition preferred phase distribution is bimodal with high probability for units to be entrained to either the peak or trough of the LFP beta cycle. Dopamine down-modulation by haloperidol shifts pallidal preferred phase to the peak of the beta cycle, while dopamine up-modulation by apomorphine and amphetamine shifts pallidal preferred phase to the descending phase (Apo1) or trough (amphetamine) of the beta cycle.

Given the opposing effect of dopamine up- and down-regulation on beta oscillations frequency, we decided to further examine the effect of beta-frequency on the preferred phase of entrained units. For that purpose, we took only units recorded in the control condition (saline) that were both oscillatory and entrained to the LFP beta cycle (Fig.S9). Units were segregated into low and high beta-frequency clusters according to their beta oscillation frequency with the cutoff at 15Hz. In pallidal units, low-frequency beta oscillations increased the entrainment to LFP beta cycle peak and decreased the entrainment to LFP beta-trough (Fig.5C middle, P=3.88, p=0.049). We then repeated this analysis for all the oscillatory entrained units included in this study, regardless of their drug condition, with similar results (Fig.5C right, P=11.91, p=0.0006). We also conducted a similar analysis using the LFP beta-frequency as the grouping factor instead of the unit beta-frequency. This analysis did not reveal any significant effect when applied on saline units. However, when applied on the entire cohort it revealed a significant effect for all unit types. Low-frequency LFP beta oscillations increased the entrainment of units to LFP beta-peak and decreased the entrainment to LFP beta-trough (wide: P=8.48, p=0.004; narrow: P=13.87, p=0.0002; pallidal: P=12.51, p=0.0004). As expected, in all cell-type categories, low-beta clusters were heavily occupied by low dopamine haloperidol recordings and high-beta clusters were mostly composed of high dopamine amphetamine and Apo1 recordings. Control and Apo2 recordings distributed equally between the two groups.

### In human PD patients, frequency of beta oscillation is modulated by DRT and disease progression

In humans, beta range is wider than in NHP, and is divided into two bands: low-beta (13-23Hz) and high-beta (23-35Hz) (Fig.6A). A PD patient can exhibit beta oscillations in either or both beta ranges, as can be seen in the single-peak or double-peak shape of the PSD (Fig.6A). In order to account for these differences we constructed a separate mixed linear effect model (MLEM) for low-beta and high-beta oscillations (Table S9). We used MLEM to estimate the effects of DRT and time on frequency and power. Since PD is a progressive neurodegenerative disease, the time factor probably coincides with the disease-induced chronic degeneration of dopaminergic cells and decrease in dopamine tone. Therefore and given our NHP results, we hypothesized that in PD patients we will see a progressive decline in beta-frequency with time. Our dataset is both nested, i.e. each patient has several recording sites with several observations per site, and unbalanced, i.e. there are different number of recordings per patient. MLEMs can be used on nested and unbalanced datasets and also overcome patient-specific variabilities.

**Fig 6.**
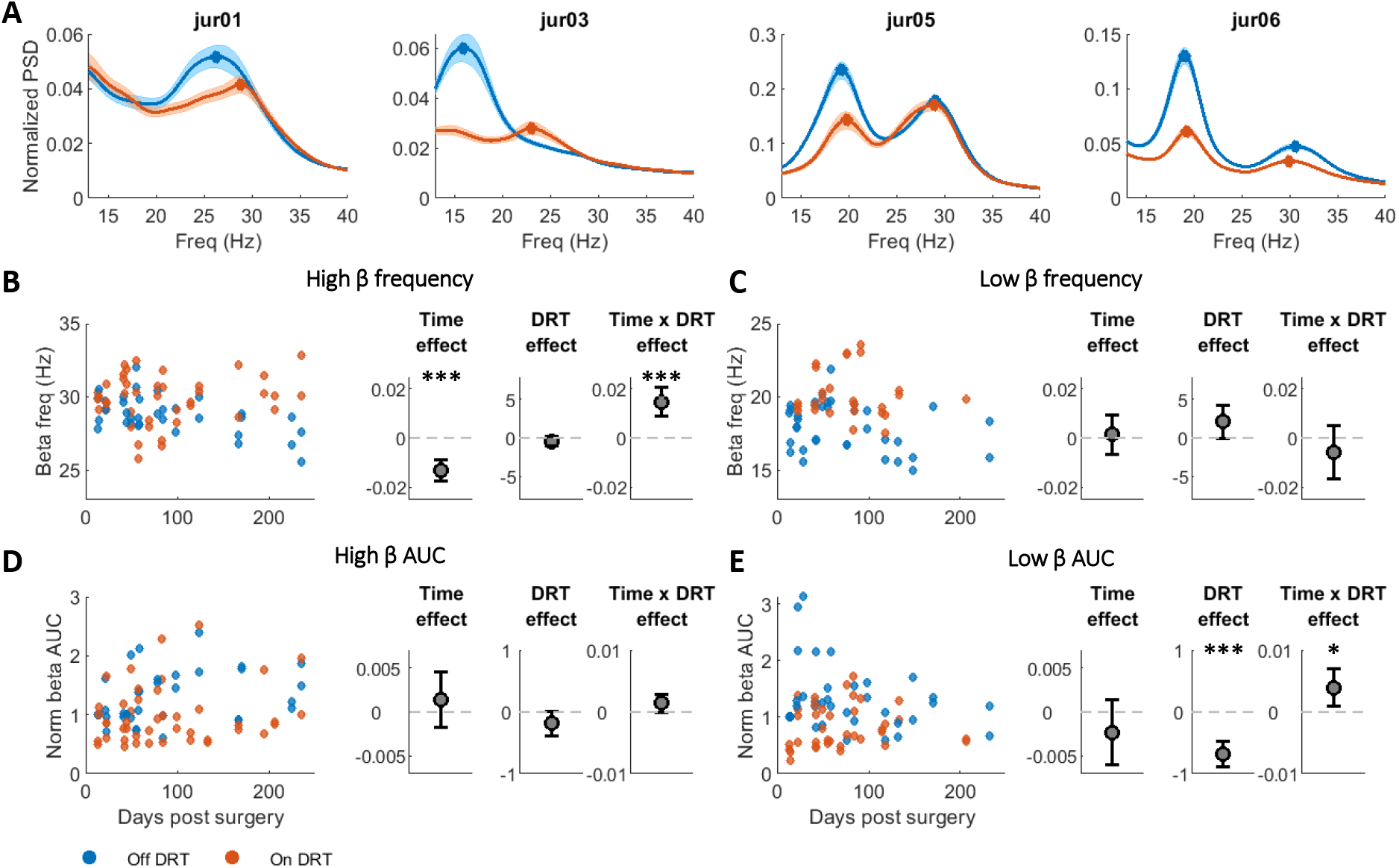
Dopamine modulation shifts LFP beta frequency in human PD patients. (A) Average nPSD off (blue) and on (red) DRT in each patient. (B-C) Time and DRT effects on beta frequency in the high (B) and low (C) beta domains. (Left) Frequency of beta peak in the high/low beta domains as a function of time post-surgery. Each point represents average per day of beta peak frequency in one STN on (red) and off (blue) DRT conditions. (Right) Time, DRT, and interaction effects on beta frequency estimated by a mixed linear effect model (MLEM). Gray points indicate each factor’s coefficient in the MLEM. Positive and negative coefficients indicate positive and negative linear relation, respectively. In the interaction effect, the coefficient presented is of time given on DRT condition. Whiskers indicate the confidence interval. The significance of the fixed effects was estimated with ANOVA test (Table S8). (D-E) Time and DRT effects on beta power in the high (D) and low (E) beta domains. Plot conventions same as (B-C). Beta power evaluated as baseline-corrected area under the curve (AUC) of normalized PSD (nPSD) in the high (D) and low (E) beta domains.

Assessment of beta oscillation frequency over a period of 250 days post-surgery exposed a consistent decline in frequency of high, but not low, beta oscillations (F(1,462)=33.95, p=1.1e-8). This decline was more robust in the off-DRT condition relative to the on-DRT condition (interaction effect: F(1,462)=25.36, p=6.8e-7) (Fig.6B). DRT did not significantly affect beta-frequency. However, in the low-beta range DRT-induced an up-shift in beta-frequency that was close to significance (F(1,414)=3.74, p=0.054) (Fig.6C). Furthermore, a visual inspection suggested a shift-up of beta-frequency post DRT in some, but not all, of the patients (see jur01 and jur03 in Fig.S10).

The effect of dopamine tone modulation by DRT generated similar trends on power in high and low-beta domains. However, the model indicated that DRT-induced decline in power was significant only in the low-beta domain (F(1,665)=40.93, p=2.9e-10) and close to significance in the high-beta domain (F(1,736)=3.35, p=0.068). This can also be seen on the single patient level (Fig.6A, Fig.S11, Table S10). DRT affected the time-induced changes in low-beta-power (interaction effect: F(1,665)=6.39, p=0.012) suggesting that off-DRT beta-power was reduced over time, while on-DRT beta-power was mildly elevated (Fig.6E).

### Up- and down-modulation of dopamine tone shifts the frequency of beta coherence in PD patients

In PD patients LFP-LFP coherence frequency was mildly dependent on dopamine tone (Fig.7A). DRT didn’t have a significant effect on the frequency of beta coherence in either high or low-beta domains (Fig.7B,C, full model results in Table S10). Chronic dopamine degeneration that induced a slow decline in beta-frequency within the high-beta domain (Fig.6) also induced a decrease in frequency of high-beta coherence (Fig.7B, F(1,120)=8.06, p=0.005), but not low-beta coherence (Fig.7C; F(1,201)=0.55, p=0.46). This can also be seen at a single patient level (Fig.S12). The model did not reveal any significant medication or time-dependent changes in LFP synchrony (Fig.7D,E). On the single patient level, the effects of time on synchrony in the high and low-beta domains are inconsistent between the patients (Fig.S13).

**Fig 7.**
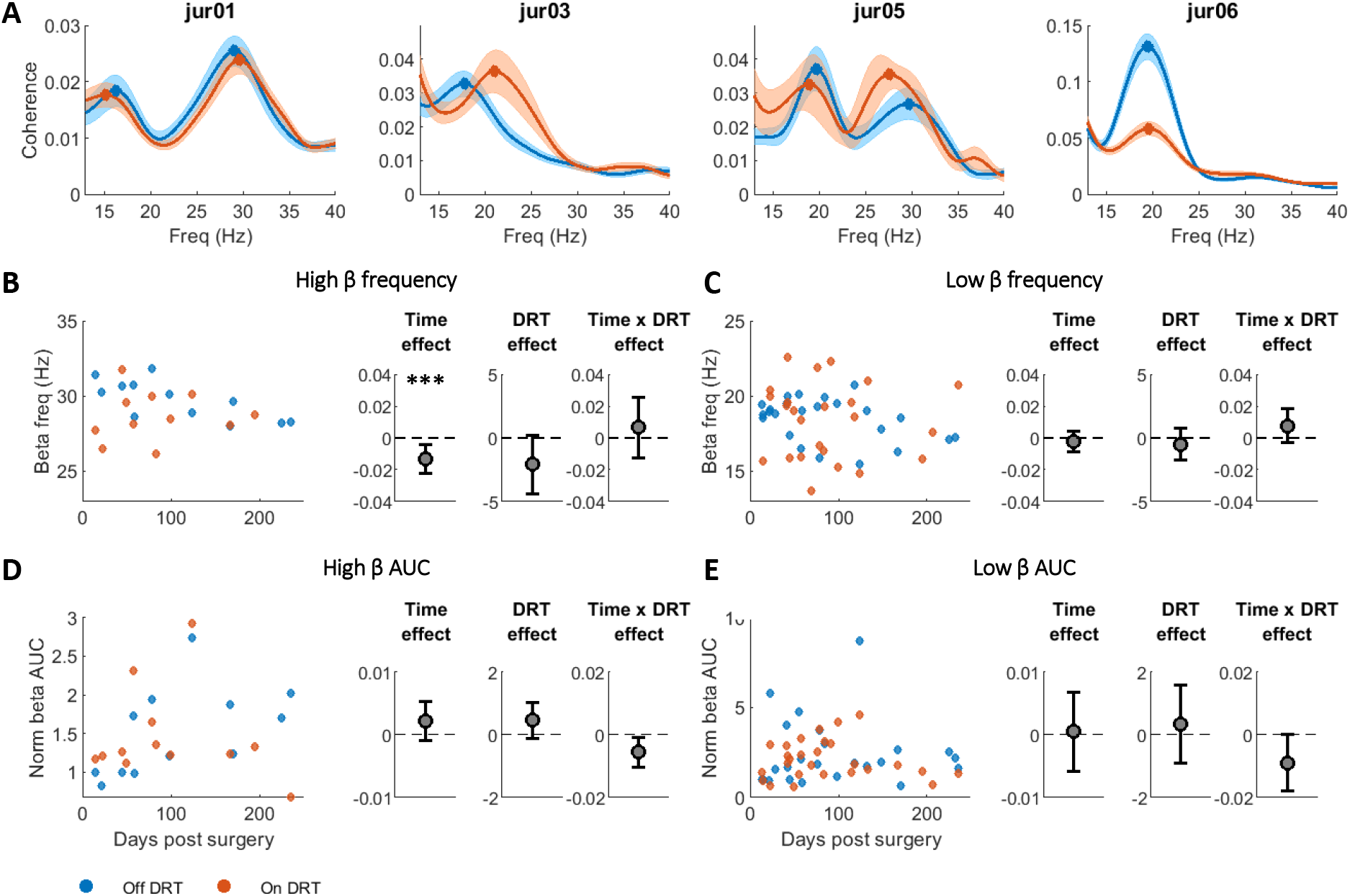
Dopamine modulation shifts LFP coherence beta frequency in human PD patients. (A) Average coherence off (blue) and on (red) DRT in each patient. (B-C) Time and DRT effects on beta coherence frequency in the high (B) and low (C) beta domains. (Left) Frequency of beta coherence peak in the high/low beta domains as a function of time post-surgery. Each point represents average per day of beta coherence peak frequency on (red) and off (blue) DRT conditions. (Right) Time, DRT, and interaction effects on beta coherence peak frequency estimated by a mixed linear effect model (MLEM). Gray points indicate each factor’s coefficient in the model. Positive and negative coefficients indicate positive and negative linear relation, respectively. In the interaction effect, the coefficient presented is of time given on DRT condition. Whiskers indicate the confidence interval. Significance of the fixed effects was estimated with ANOVA test (Table S8). (D-E) Time and DRT effects on beta synchrony in the high (D) and low (E) beta domains. Plot conventions same as (B-C). Beta synchrony is evaluated as baseline-corrected area under the curve (AUC) of the coherence in the high (D) and low (E) beta domains.

## Discussion

Analysis of beta properties in NHP and PD patients showed that the frequency of beta oscillations and beta coherence strongly correlated with acute and chronic changes in dopamine tone. In contrast, effects of dopamine tone on beta-power, beta synchrony and spike/LFP phase-locking are non-monotonic, inconsistent and less robust (Fig.8). These results emphasize beta-frequency, and not beta-power or any other beta oscillation features, as the key property of physiological and pathological beta oscillations in CBG networks.

**Figure 8:**
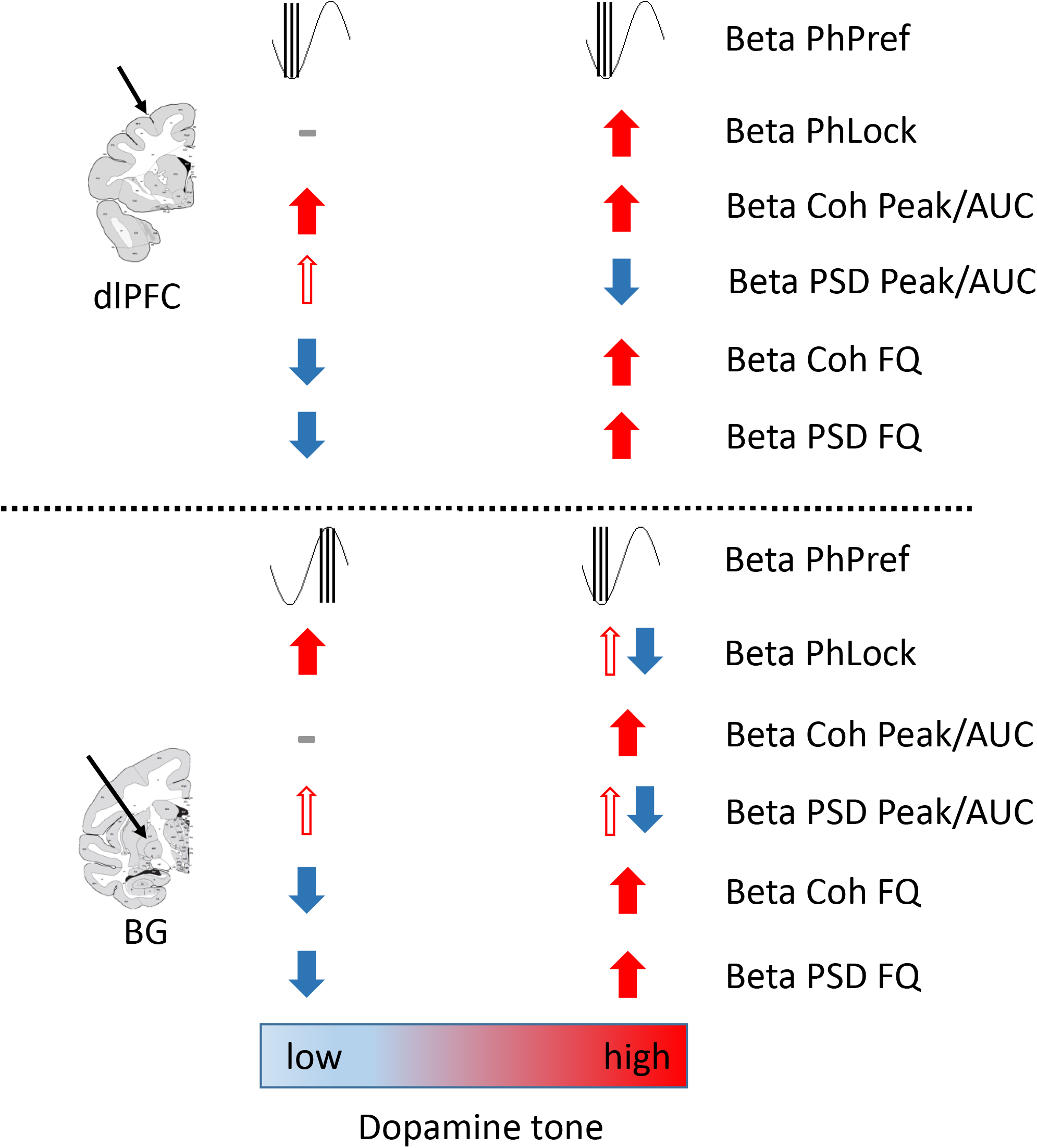
Results Summary. Synopsis of LFP and SUA results. Thick arrows indicate statistically significant effects. Thin arrows indicate trends that did not reach statistical significance. Beta PhPref: spikes preferred phase in LFP beta cycle; Beta PhLock: spike to beta LFP phase locking; Beta Coh Peak/AUC: beta synchrony, measured as coherence beta peak/AUC; Beta PSD Peak/AUC: power of beta oscillation, measured as nPSD beta peak/AUC; Beta Coh FQ: frequency of beta coherence; Beta PSD FQ: frequency of beta oscillation. In the BG Beta PSD Peak/AUC sections thin red lines represent trends that were significant for the top 20% of all units.

### Beta-frequency but not beta-power is monotonically correlated with acute changes in dopamine tone in PD patients and NHPs

Previous studies in PD patients and animal models have examined the effects of acute dopamine tone changes by assessing properties of beta oscillations during off and on DRT. These studies suggested that while STN LFP beta-frequency is not altered by acute changes in dopamine tone^11,13,23–25^, beta-power is decreased by acute increase in dopamine tone following treatment with L-Dopa and apomorphine^13,23,25–27^. In our study, acute dopamine up- and down-modulation in NHP resulted in clear shifts up and down the beta oscillation frequency domain (Fig.2-3). Remarkably, beta-power increases could be generated in the LFP (Fig.2) and SUA (Fig.3) by either acute up- or down-regulation of dopamine tone. These results suggest that in healthy systems both up- and down-modulations of dopamine tone have the capability to generate high-power beta signals. Regarding the opposite effect of dopamine up-modulation by amphetamine and apomorphine (Fig.2) on beta-power, we can assume that this effect depends on additional parameters other than purely dopamine tone.

Both amphetamine and apomorphine increase dopamine neurotransmission tone, but mechanisms and sites of action are fundamentally different between these two agents. Amphetamine inhibits DAT function, which leads to increased activity-dependent dopamine concentration in the synaptic cleft and stimulation of synaptic dopamine receptors. Previous studies in rodents and humans measuring beta activity after DAT inhibition, via administration of methylphenidate and cocaine, showed a sharp increase in beta-power^28,29^. Apomorphine, on the other hand, directly stimulates dopamine receptors, probably with higher affinity for D2 receptors, in an activity-independent manner and irrespective of their location^30^. Consequently, apomorphine activates not only synaptic and extra-synaptic receptors in the projection nuclei but also autoreceptors that are activated by the somatodendritically released dopamine. Once autoreceptors are activated, dopamine dynamics can be altered via inhibition of the discharge of dopamine neurons, as well as by reduced dopamine synthesis and release^31–33^. These and other differences between amphetamine and apomorphine could be behind the variability in dynamics of beta-power response.

Our results of LFP beta oscillations in PD patients off/on DRT (Fig.6) are in-line with previously published reports confirming a significant medication effect on lowering of beta-power. Here, we have shown that the effect of medication on beta-power is accompanied by its effect on beta-frequency. Hints of this phenomenon can be found in previous reports^13,25^.

### Beta-frequency but not beta-power is correlated with chronic changes in dopamine tone in PD patients

Previous studies of beta properties after chronic dopamine modulation in human and animal models produced inconsistent results^11,26,34^. In rodents, chronic dopamine denervation resulted in either no change in frequency or power of beta^26,34^, or a progressive increase in beta-power but no change in beta-frequency^10^. In primates, beta-frequency was shown to decline with progressive parkinsonism caused by staggered MPTP administrations^35,36^. This decrease in beta-frequency was accompanied with an increase in power. A human study assessing beta-frequency over a 7 year period in post-surgery PD patients showed no change in beta-frequency and a decrease in beta-power^37^. Our results showed a consistent and significant time effect post-surgery in high beta-frequency, but beta-power changes were less consistent (Fig.6B-D, Fig.S10,11). The difference in methods of data collection could explain the disparity between our results and those previously published^37^. Our method involved multiple recordings in the off and on states from each of the 4 patients over a span of 170 to 250 days post-surgery, creating a chronological trajectory for each patient off and on DRT. While dopamine tone was not directly measured in our patients, persistent dopaminergic degeneration has long been established in PD. These results further support our hypothesis that progressive beta-frequency decline, and not a beta-power amplitude, could be used as a more effective marker for the progression of PD and as a trigger for adaptive DBS procedures

### Coherence frequency within the beta domain but not degree of synchronization is monotonically correlated to acute and chronic changes in dopamine tone

Multiple reports concluded that coherence in the beta range within and between the CBG networks is elevated in a parkinsonian state^6,10,12,15,17^. However, the effect of acute dopamine modulation on beta-range coherence is still debated. Dopamine-depleted rodents showed a decrease in coherence in the beta range after apomorphine administration^12^, accompanied with an increase in coherence frequency. Conversely, PD patients off and on DRT showed no change in beta coherence^23^. Here, we showed that acute changes in dopamine tone in NHPs shifted the frequency of LFP coherence in the beta domain within and between dlPFC and GPe (Fig.4). These shifts mimicked the shifts in beta-frequency in the power spectrums of single sites/units (Fig.2 and 3). In PD patients, presumed chronic dopamine degeneration resulted in consistent decline in coherence frequency within the high-beta-frequency domain (Fig.7). However, acute changes in dopamine tone did not significantly affected the degree of synchronization. We suggest that it is not sufficient to rely on the degree of beta coherence when studying dopamine modulation effect on the CBG circuit. Attention should be paid to the frequency features of coherence.

### Phase of spike-LFP entrainment is differentially affected by acute changes in dopamine tone

Entrainment of spikes to a specific phase of LFP oscillation is a ubiquitous phenomenon in neural circuits. Previously published analysis supports the notion of an increase in single-unit phase-locking to beta LFP with dopamine loss^38^. In the GPe, haloperidol-induced entrainment was higher than in control as evidenced in the number of entrained units and degree of entrainment. When we further examined the relationship between beta-frequency and spike phase preference, we found that low-beta-frequency entrained spikes to the peak of LFP beta oscillation while high-beta-frequency preferentially locked spikes to the trough of the LFP beta cycle (Fig.7D). This suggests a relationship between beta-frequency and phase preference, and that in addition to the frequency of beta oscillation, the preferred phase of spike to LFP locking could also be a factor in the behavioral outcomes. Previously published reports show that inputs arriving at the optimally excitable phase of beta cycle of cortical cells lead to responses that are greater in amplitude and have a shorter latency of motor evoked potentials^39^. Additionally, this phase-dependence was found to be more robust during low-beta (16-19Hz) as opposed to high-beta expression^40^.

## Conclusions

Dopamine transmission pathologies lie at the root of many neurological and psychiatric diseases, like PD, depression, schizophrenia, and ADD/ADHD. In dopamine-depleted PD patients and animal models increase in beta-power has become a hallmark of the disease. However, high beta-power was also found in the CBG of subjects with no movement disorders^41,42^. Our study demonstrated that beta-frequency is a more reliable and accurate marker of acute and chronic up- and-down regulation of dopamine tone. Changes in beta-frequency were detected after acute modulation in dopamine tone in NHP and during chronic progression of PD in human subjects. Coherence frequency mimicked shifts seen in beta-frequency. Whereas previous studies proposed that high beta-power is an indication of dopamine degeneration, our data showed that increases in beta-power can be generated via bidirectional shifts in dopamine tone. Finally, acute up- and down-modulation of dopamine tone can lock spikes to opposite phases of beta LFP. Thus opening the possibility that the spike preferred phase, along with oscillation frequency, contribute to the electrophysiology behind the PD akinetic phenotype.

Further studies that can simultaneously record neuronal activity and detect extracellular dopamine levels could be useful in elaborating the relationship between dopamine tone and beta oscillation frequency. To better understand the unique patterns of oscillatory activity between the normal, hypo- and hyper-dopaminergic states it would be critical to decipher the relationship between beta oscillations and other frequency bands. Our study is limited because we did not selectively affect particular dopamine targets in a precise nucleus within the CBG network, or modulated specific dopaminergic circuits, or selectively stimulated pre-versus post-synaptic dopaminergic receptors. Future studies should further investigate the relationship between dopamine tone and beta activity. A better understanding of this unique physiological phenomenon can improve patient care, impact the therapeutic potential of aDBS procedures, and can serve as a foundation and a springboard for further clinically-relevant exploration.

## Star Methods

### Contact for Reagent and Resource Sharing

Further information and requests may be directed to and will be fulfilled by the Lead Contact, Dr. Lily Iskhakova

### NHP experiment: Experimental Model and Subject Details

All experimental protocols were conducted in accordance with the National Institutes of Health *Guide for the Care and Use of Laboratory Animals* (National Research Council, 2011) and with the Hebrew University guidelines for the use and care of laboratory animals in research. The experiments were supervised by the Institutional Animal Care and Use Committee of the Faculty of Medicine, the Hebrew University. The Hebrew University is an Association for Assessment and Accreditation of Laboratory Animal Care internationally accredited institute.

Data was collected from two healthy, young-adult, female vervet monkeys (*Chlorocebus aethiops sabaeus*; G (MD-13-13518-4) and D (MD-15-14412-5)) weighing ~4 kg. Monkeys were obtained from the St. Kitts Monkey Farm. The age of monkeys at the time of the experiment was 5-8 years (G: 7-8, D: 5-6). Each monkey was trained in the task ~3 months prior to the implantation surgery and recording. After completion of recordings the chambers were removed. Once the monkeys recovered, they were both transferred to the Ben Shemen Israeli Primate Sanctuary (www.ipsf.org.il/).

### Surgery and Post-Operative Maintenance

Recording chamber implantation took place after the monkeys were fully trained in the task (~ 3 months, 5-6 days a week, 500-1500 trials per day). An MRI-compatible Cilux head holder (Crist Instruments, MD) and a rectangular 34×27 mm (inner edge) Cilux recording chamber (AlphaOmega Engineering, Israel) were implanted above a burr hole in the skull under deep anesthesia in aseptic conditions as described previously^5^. The head holder and the chamber were attached to the skull using titanium screws (Crist Instruments, MD) and wires (Fort Wayne metals, IN) embedded in acrylic cement. The central and arcuate sulci of both left and right lobes were within the limits of the chamber, which provided bilateral access to the dlPFC and GPe (Fig.1A,B). Finally, two titanium ground screws (Crist Instruments, MD) were placed in contact with the dura mater and connected to the chamber and head holder using a titanium wire.

Recordings began after a postoperative recovery period of 7 to 10 days, during which an anatomical MRI scan was performed^5^ to estimate the chamber coordinates of the neuronal targets. Throughout the entire course of the experiments, the chamber was washed with saline solution every 24-48 hours. After the completion of the recordings an MRI scan was performed to confirm the location of the recording sites and to rule out significant brain shifts.

### Task and behavior monitoring

NHP subjects were engaged in a behavioral task. The analysis of task behavior-related electrophysiology is out of the scope of the current publication. Briefly, a reversal-learning task included multiple blocks with trials consisting visual cues predicting outcomes of different valences. Task included five types of outcomes: palatable and less-palatable food, air puff to the eyes or nose, and neutral outcome (no food or air puff delivered). Predictive cues were shuffled between blocks, so the animals had to learn the new cue-outcome mapping at each block. Anticipatory and event-related behavior was recorded via laser lick sensors (Sick Sensor Intelligence) and eye-tracking device (ISCAN Incorporated).

### Acute Dopamine Modulation

All injections were made during the last trial of the first block (20-25 minutes after the beginning of the recording, Fig.1C). Saline (0.1cc), haloperidol (1 mg/kg) and amphetamine (0.5 mg/kg) were injected intramuscularly (IM) and apomorphine (0.5 mg/kg) was injected subcutaneously (SubQ). All injections were performed in accordance with the manufacturer instructions. For analysis purposes, drug influence time was considered to begin five minutes after drug injection and lasts until the end of the task (>3 hours). Apomorphine has a fast dynamic. Its initial effect lasts for ~1 hour and then activity returns to baseline. Therefore, we divided post-apomorphine recordings into two phases, agonistic phase 5-60 minutes after the injection (Apo1) followed by a post-agonistic phase that lasted until the end of the task (Apo2).

### *In vivo* electrophysiology

During the recording sessions, the monkeys’ heads were immobilized with a head-holder. Local field potentials (LFPs) and single-unit spikes were simultaneously recorded (Fig.1D) from eight glass-coated tungsten electrodes (impedance 0.45-0.8 MΩ measured at 1000Hz). Electrodes were arranged in two towers with four electrodes per tower. Each tower was localized to allow targeting and recording from three configurations: bilateral GPe or dlPFC, and unilateral (left or right) GPe/dlPFC. Electrodes were navigated within the brain using the Electrode Positioning System (AlphaOmega Engineering, Israel).

The electrical activity was amplified by 5000, high-pass filtered at 1Hz using a hardware two-pole Butterworth filter and low-pass filtered at 10 kHz using a hardware three-pole Butterworth filter. Raw data was sampled at 44 kHz by a 16-bit (± 1.25V input range) Analog/Digital (A/D) converter. LFP was low pass filtered at 200 Hz and sampled at 1375Hz. (AlphaLab SnR Stimulation and Recording System, AlphaOmega Engineering, Israel)

Spiking activity was sorted online using a template matching algorithm. Up to four different units could be simultaneously isolated from the same electrode. Off-line, the isolation quality of each unit was graded by calculating its isolation score^43^. The isolation score ranged from 0 (i.e., multi-unit activity) to 1 (i.e., perfect isolation). Only units that were recorded for over one minute and had isolation score ≥0.7 were included in the database. Description of the full dataset can be found in Table S1.

### Unit type identification

GPe units were identified as either high frequency discharge (HFD) neurons or low-frequency discharge (LFD) neurons based on their firing rate and pattern of activity^44^. Units with firing rate above 30 Hz were classified as HFD. Units with firing rate below 30 Hz were manually identified as either HFD’s or LFD based on absence or presence of bursts, respectively.

Cortical units were separated into putative-pyramidal cells and putative-interneurons based on the width of the spike shape^22,45^. The separation criteria was determined according to spike width distribution (Fig.S1). Units with spike width larger than 3 SD over the mean were excluded from further analysis.

### Spectral analysis

#### Power spectrum density (PSD)

Power spectrum density (PSD) of LFPs and single-units was calculated using the welch method. For LFPs, the signal was first cleaned of high-amplitude artifacts, defined as deviation of over 5 SD from the signal mean. Once such deviation was detected, the surrounding points were also included in the artifact until the LFP resumed value within 3 SD from the mean. Artifacts were replaced with zeros, which do not influence spectral analysis results. The clean LFP was parsed into one-minute segments with 54 second overlap. For each time (see above).

#### Identification of oscillatory signals

To classify either LFP sites or single-units as oscillatory, we examined their nPSDs and looked for a peak in the beta range (8-24 Hz). We required that the prominence of the beta-peak (a measurement of peak size relative to its surrounding) would be sufficiently larger than the noise-level peak-prominence of the nPSD. Noise level was estimated as the median of the peak-to-trough distance in the nPSD. A beta-peak that was two times larger than the noise level was considered sufficiently large, and the LFP/ single-unit was classified as oscillatory (Fig.S3,14, Table S5). Effect of drug injection on percentage of oscillatory sites/units in the post-drug conditions was tested using chi-square test followed by post-hoc comparison with Bonferroni correction for multiple comparisons (Fig.S3,14, Table S5).

#### Beta oscillation properties

Total beta-power was estimated in two ways: (1) As the nPSD’s area under curve (AUC) in the beta range, (2) as the peak value in the beta range. These methods were selected to overcome each other shortcomings. The peak value is a direct estimation of beta power in its most prominent frequency, but it is affected by the typical 1/f shape of the nPSD. i.e. peak values in lower frequencies tend to be larger. The AUC measures the total power in the beta band and therefore it is a more general estimation that is less affected by the location of the peak. If no significant peak was found (i.e. non-oscillatory LFP/unit, see above), mean nPSD value in the beta range was utilized instead. Mean nPSD value was preferred to maximum value because the later tends to detect power at the edges of the beta band and reflects the nPSD shape rather than the beta-power per se. For oscillatory signals we also extracted the beta band peak frequency. If more than one peak was found within the beta range, the peak with the highest prominence was chosen for the beta peak and frequency analysis. Signals with beta-AUC larger than 5 SD above the mean were defined as outliers and excluded from further analysis. To test for significant effect of dopamine modulation on beta properties (beta AUC, peak and frequency) we preformed Kruskal-Wallis test, followed by post-hoc Tukey test. Here and in other analyses below Kruskal-Wallis test was chosen as a non-parametric alternative for ANOVA since our data failed to fulfill the test assumptions. Analysis was conducted with matlab built-in functions. In the Tukey post-hoc test, matlab function (multcompare) did not deliver p values smaller than 10^−10^ due to parameters involving the estimation of the studentized range cumulative distribution function (CDF). We chose to use the built-in parameters since their accuracy is more than enough for statistical inference. Beta properties of LFP and single-units in the saline condition were also compared to that of the naïve animals using independent two-sample t-test, to test for a possible effect of the injection and recording procedure.

#### Coherence and phase-locking value

For each simultaneously recorded LFP signals or single-units, traces were segmented into one minute segments with 30 seconds overlap, and magnitude-squared coherence was calculated for each segment using the Welch’s overlapped averaged periodogram method with 5 seconds window, 2.5 second overlap, and frequency resolution of 1/3 Hz in 1-200 Hz range.

Since coherence between two signals is affected by both oscillation amplitude and phase synchrony^46,47^, we further calculated the phase-locking value (PLV)^46^ between each pair of signals to directly measure the later. PLV was calculated after similar segmentation in the time domain as coherence, and with 1 Hz resolution in the range of 1-80 Hz.

#### Identification of synchronized pairs

Synchronized pairs were defined using a similar method to that described above for oscillatory LFPs or single-units. Briefly, we required that the beta-peak prominence in the coherence function have signal to noise ratio (SNR) of 2 or above.

#### Beta coherence and phase-locking value (PLV) properties

As for LFP and single-unit nPSD beta properties, we assessed beta synchronization in two ways: (1) as the coherence AUC in the beta range, and (2) as the peak value in beta range or mean value if no significantly large peak was found. For synchronized pairs we also extracted the coherence beta-peak frequency. We repeated the same analysis for PLV. To test for significant effect of dopamine modulation on beta properties we preformed Kruskal-Wallis test, followed by post-hoc Tukey test.

#### Spike-to-LFP entrainment

If an LFP site was classified as oscillatory, we further analyzed the entrainment of the spike discharge of local units to the LFP oscillations. LFP was band-pass filtered around its central beta-frequency (defined above) ±2 Hz using a four-pole Butterworth filter, and beta phase was extracted using Hilbert transformation. For each unit, beta phase at the time of each spike was extracted. The degree of unit-to-LFP entrainment can be measured by the vector-length of the circular average of spike phases. Vector length values range from 0 to 1. A large vector-length indicates a high tendency of spikes to be clustered around a specific phase of the LFP oscillations. To evaluate dopamine modulation effect on vector-length we preformed Kruskal-Wallis test, followed by post-hoc Tukey test. We further evaluated dopamine modulation effect on entrained unit preferred phase. To classify units as entrained we preformed Rayleigh test for each unit followed by FDR correction for multiple comparison and used a conservative threshold of p value = 0.0001. This threshold was chosen to minimize the effect of false detection on our results. Still the entrainment analysis was more sensitive than the oscillation analysis described above (see Fig.S9). The preferred phase of an entrained unit was defined as the circular mean of its spike-phase distribution. Dopamine modulation effect on phase preference was assessed by circular median test followed by post-hoc pairwise comparison with Bonferroni correction for multiple comparisons. The circular median test was chosen over the common Watson-Williams since our data did not fulfill the latter requirements.

We further divided the oscillatory entrained units (see Fig.S9A, second column) into low-beta and high-beta groups according to the unit central beta-frequency (below and above 15 Hz, respectively). We repeated the statistical test to assess the effect of the unit beta-frequency on entrained units’ preferred phase, for units in the saline condition and for all units regardless of their drug condition. We also repeated the same analysis for all entrained units with LFP beta-frequency as the grouping factor instead of unit beta-frequency.

### Human Data

Methods for human data collection and analysis were thoroughly described in our previous publication^41^.

### Patient selection

In this study, four PD patients underwent STN DBS surgery with implantation of the Activa PC+S pacemaker (Medtronic, Inc, Minneapolis, MN, USA). All patients met accepted inclusion criteria for DBS surgery and signed informed consent. Patients had (i) advanced idiopathic PD; (ii) long-term levodopa use with reduced efficacy, on-off motor fluctuations and increased incidence of medication-induced side effects; (iii) normal cognitive function or mild-moderate cognitive decline as defined by Addenbrooke’s cognitive examination (ACE) >75 and frontal assessment battery (FAB) >10. Patients’ levodopa equivalent dose (LED) was calculated according to Tomilson et al^48^. Patient demographic and clinical information is detailed in Table S10. The study was authorized and supervised by the IRB of Hadassah Medical Center (no. 0403-13-HMO) and the Israel Ministry of Health (no. HT6752). Clinical Trials Registration number: NCT01962194.

### Intra-Operative Procedure

The surgical technique is described elsewhere^41,49,50^. Briefly, STN target coordinates were chosen using Framelink 5 or Cranial software (Medtronic, Minneapolis, USA). STN entry and exit were verified intraoperatively by microelectrode recording of multi-unit spiking activity along the trajectory. The final localization of the permanent DBS electrode was determined according to (1) analysis of spontaneous spiking activity, (2) response of spiking activity to passive and active movements and (3) clinical effects of stimulation at the target. In one of our patients (jur05), one contact was malfunctioning, and data from this contact were excluded from the analysis.

The permanent lead implanted during the surgery (model 3389; Medtronic, Inc., Minneapolis, MN) had four contacts. Each contact had a diameter of 1.27 mm and length of 1.5 mm spaced by 0.5 mm intervals. The lead was placed along the dorsal/lateral/anterior-ventral/medial/posterior axis of the STN (Fig.1E), and contacts were numbered from 0 (ventral) to 3 (dorsal). Generally, contact 1 was placed dorsally to the border between the motor dorsal beta-oscillatory region and the non-motor ventral non-oscillatory region, detected automatically by a hidden Markov model (HMM)^49^ (Fig.1F).

### Post-operative clinical assessment and electrophysiological recordings

Patients underwent recordings during 170-400 post-operative days. Recordings from the first week post-surgery were excluded from the dataset to avoid insertion effect. Only recordings from the first 250 days were included in the analysis because recordings after 250^th^ day all came from a single patient (jur01, Fig.S10-13). Post-operative recordings were acquired in an outpatient setting. Patients had clinical evaluations and recording sessions every 2–4 weeks. During recordings, patients were instructed to sit quietly for the rest-state session, which lasted three minutes. In addition, sessions included recordings during performance of four tasks, which are out of the current paper scope (Provocative Images task, Doubt task, Auditory Go-NoGo task, Emotional Voice Recognition task as described by Rappel et. al.^41^).

Recordings took place during an off-medication and on-medication states (Fig.1G,H, see Table S2 for number of recording days and sessions). Off-medication recordings took place after overnight withdrawal of DRT. On-medication recordings took place after confirmation of a substantial improvement in the parkinsonian motor clinical symptoms by the patient and the examiner.

LFP activity was recorded from all the bipolar contact pair combinations (0-1,0-2,0-3,1-2,1-3, 2-3) in both hemispheres through the Medtronic PC+S recording setting. Each contact pair recording lasted 30 seconds. The signal was amplified by 2000, band-passed from 0.5 to 100 Hz, using a 3 pole Butterworth filter, and sampled at 422 Hz by a 10-bit A/D converter (using ± 2V input range).

### Spectral analysis

#### Signal preprocessing

LFP signal was filtered between 0.5 and 200 Hz using four-pole butterworth IIR filter. We observed three types of noise artifacts in our data: ECG artifact, transient high noise artifacts, and line noise artifact. First, we removed ECG artifacts related to the Activa PC+S device. As seen in our data^41^ and reported by Swann et al^16^, bipolar recordings from the most ventro-medial electrode contact (Contact 0, Fig.1F) were accompanied by electrocardiogram (ECG) artifact in three out of our four patients. This artifact probably originates from current leakage into the Activa PC+S at the insertion site of the device lead extender over the pectoralis muscle^16^. ECG pulses were identified by their high peak (above 1.2 SD over the mean) and regularity (coefficient of variation (CV) < 0.32). The ECG signal recorded from the bipolar contacts 0-1, 0-2, 0-3 was averaged to create a template for each bipolar recording for each visit. This template was subtracted from every occurrence of the ECG artifact using linear regression to achieve optimal fit with the data^41^. Second, we removed transient high noise artifacts from the data. Transient high noise artifacts were identified according to their large absolute amplitude (>5 SD over the mean). Noise start and end points were defined as return of the absolute amplitude to ≤ 3 SD distance from the mean. Line noise artifact was removed directly from the PSD (see below).

#### Spectral analysis

PSD was calculated using the welch method with 500 millisecond windows, 100 millisecond overlap, and frequency resolution of 1/2 Hz. PSD was divided by the total power to get the normalized PSD (nPSD). The total power was calculated as PSD sum across frequencies, excluding the PSD portion that was influenced by line noise artifact (46-57 Hz). The wide range of omitted frequencies was selected to minimize the effect of line noise artifact on the results of beta oscillation properties analysis.

#### Beta oscillation properties

In the analysis of PD patients’ LFP beta properties we took a similar approach as described above for the NHP LFP data. However, there are some differences between human and NHP characteristic beta oscillations. Human beta range spans higher frequencies, and it is common to find beta oscillations in two separate beta ranges, low (13-23 Hz) and high (23-35 Hz), in some but not all patients. Therefore, we defined active beta range (low-beta, high-beta or both) manually for each patient according to their mean nPSD. In patient jur03 there was only low-beta activity but its frequency range was exceptionally wide, so for this patient low-beta range was set to 13-25 Hz.

For each observation (recording from a single bipolar pair at a single session) we calculated beta power and frequency. Beta-power was defined as the nPSD’s AUC in the beta range. To overcome patient variability in baseline power estimation we performed baseline correction. Beta power in the first recording day at each bipolar pair was subtracted from all consecutive beta power values of the same bipolar pair. Beta-frequency was defined as peak value frequency, if a significant peak was found. Significant peaks were defined relative to other peaks in the same nPSD. We considered a beta peak to be significant if its prominence z-score was equal to or greater than 0.67, equivalent to 75^th^ percentile in a standard normal distribution.

To assess the contribution of dopamine replacement therapy (DRT) and disease progress to beta power and frequency we constructed a linear mixed effect model (MLEM). The model included fixed effects of DRT (on/off) and time from surgery and for the interaction between the DRT and time factors. The model also included random terms for intercept, DRT and time effects for each patient and each bipolar pair in each hemisphere. We constructed a separate model for beta frequency in low-beta and high-beta range, which included only significant peaks. We also constructed models for beta power in low and high beta range, which included all recordings.

#### Beta coherence properties

Magnitude-squared coherence was calculated for each segment using the Welch’s overlapped averaged periodogram method with 250 milliseconds window, 125 milliseconds overlap, and frequency resolution of 1/10 Hz in 1-100 Hz range. Beta analysis was similar to that described above for PSD. Again, beta ranges were manually assigned to each patient according to their average coherence.

## Data and Software Availability

Data analyses were performed with custom written scripts in MATLAB (Mathworks, Natick, MA). Requests for data and MATLAB scripts used in the present study can be directed to the lead author. Data will be posted on the lab website and made available upon request.

## Acknowledgments

We thank Dr. Uri Werner-Reiss for assistance with training and care of monkeys. Anatoly Shapochnikov for help designing and building our hardware and set-up. The veterinary department at Hadassah Medical Center for health maintenance of our primates. The work was supported by the grants from the Teva: National Network of Excellence (L.I.), Israeli Ministry of Absorption (L.I.), Edmond and Lily Safra Brain Center (L.I., P.R.), Rosetrees and Adelis foundation and ISF grants to HB. ISF grants (1129/12 and 2128/19) to R.E., NIPI grant to R.E,.

## Author Contributions

L.I, P.R., and R.E, H.B. conceived the research and designed the experiments. Z.I. performed the surgeries. L.I, P.R., G.F. performed the in vivo physiology experiments and data analysis. R.E. and O.M. performed the human physiology experiments and data analysis. L.I. and P.R. wrote the manuscript and all authors commented on and approved the writing.

