## Supplemental Figures for "Frequency matters: Up- and Down-Regulation of Dopamine Tone Induces Similar Frequency Shifts in Cortico-Basal Ganglia Beta Oscillations"

**Supplementary figures**

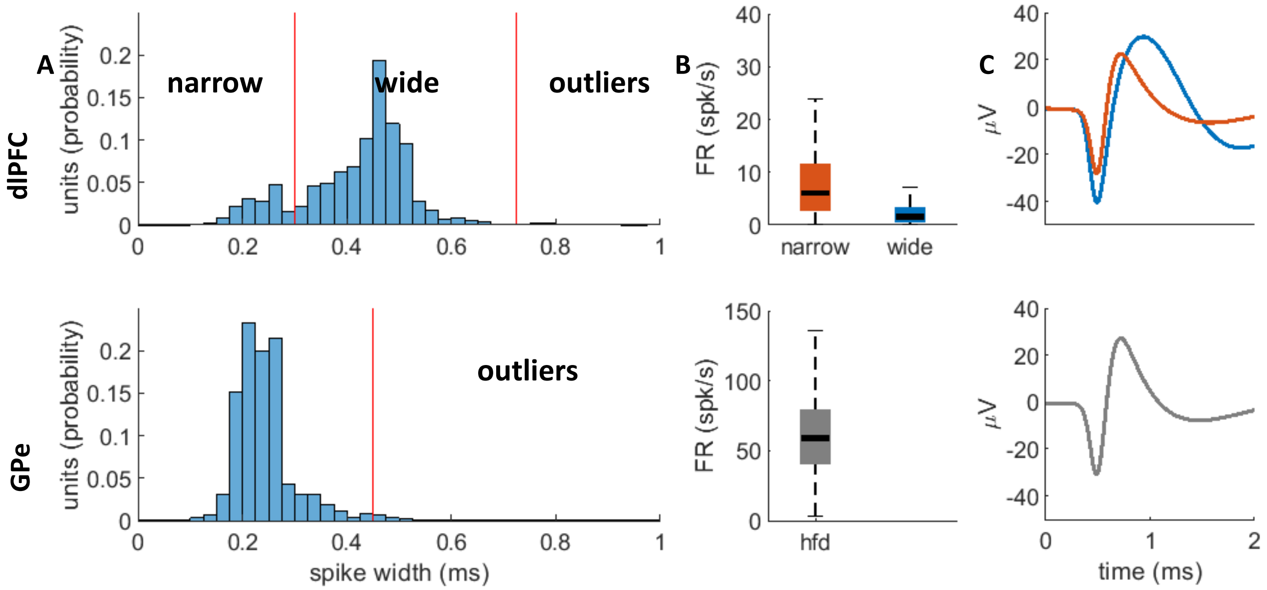


**Fig S1. Description of cortical and pallidal single unit spike shape and firing rate properties.** Top row: dlPFC units. Bottom row: GPe units. (A) Spike width histogram. Cortical wide and narrow units were defined according to their spike width (trough to peak). Units with spike width that exceeded 3 SD over the mean were considered as outliers and excluded from the dataset. (B) Single unit firing rate. On each box, the central line indicates the median, and the bottom and top edges of the box indicate the 25th and 75th percentiles, respectively. The whiskers extend to the most extreme data points within 1.5*IQR (interquartile range, equal to the length of the box) distance from the edge of the box. Narrow units had higher firing rate relative to wide units, in-line with their identification as putative interneurons and pyramidal cells, respectively. (C) Average spike shape of cortical wide (blue), narrow (orange), and pallidal HFD (gray) units.

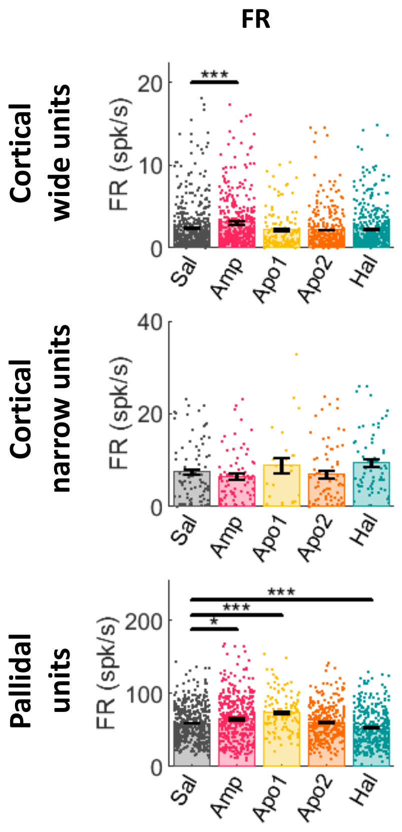


**Fig S2. Single unit FR.** Top: cortical wide units. Middle: cortical narrow units. Bottom: pallidal units. Bars indicate average values. Single points indicate individual unit values within the range of mean ± 3 SD. Black vertical lines indicate standard error of the mean. Drug influence was evaluated by Kruskal-Wallis test followed by post-hoc Tukey test. * $p<0.05$ ** $p<0.01$ *** $p<0.001$

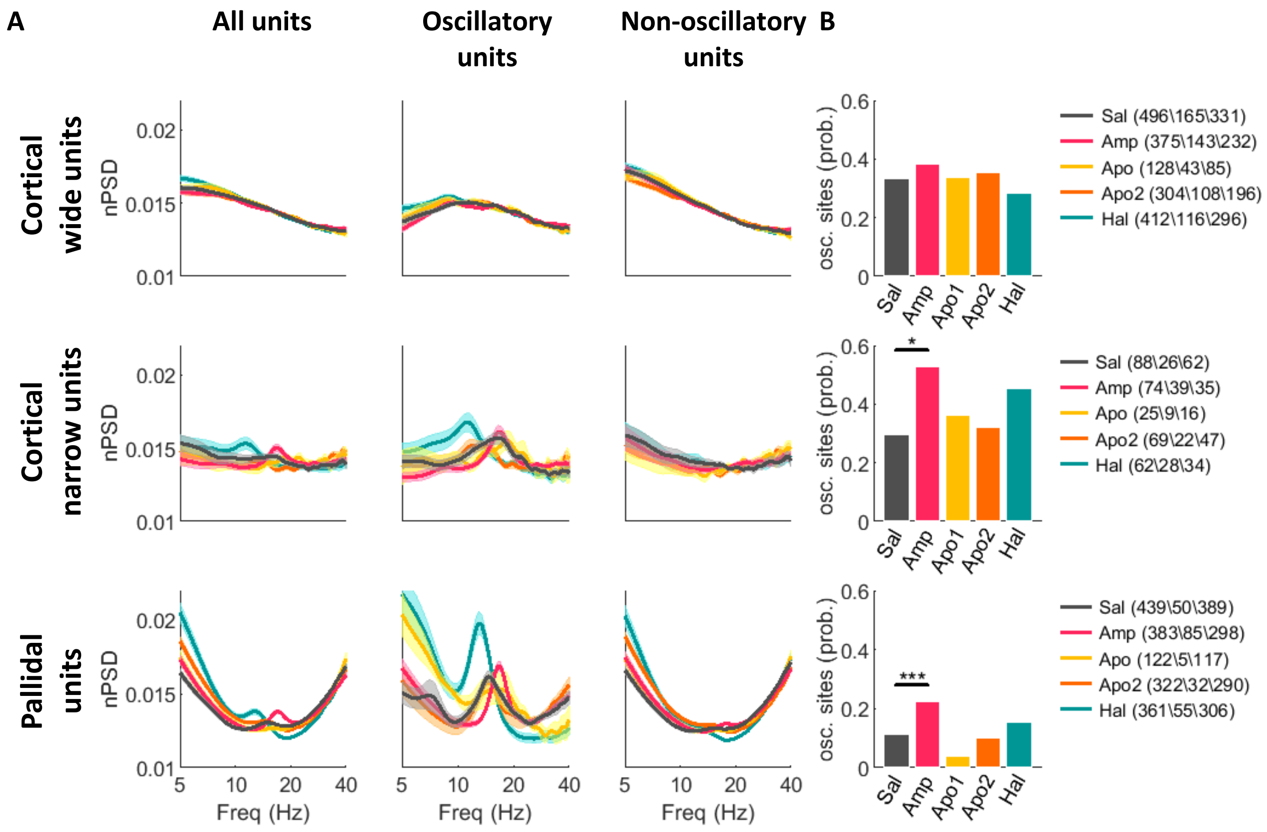


**Fig S3. Identification of oscillatory units.** (A) Average nPSD of all (left), oscillatory (middle), and non-oscillatory (right) single units in all drug conditions. (B) Probability of oscillatory units out of all the recorded units in each condition. Drug effect was tested with chi square test (wide: $\chi^2_{(4)}$=9.47, p=0.0505, narrow: $\chi^2_{(4)}$=11.70, p=0.0197, pallidal: $\chi^2_{(4)}$=38.75, p=7.8e-8) followed by pairwise comparisons with Bonferroni correction for multiple comparisons. Comparisons between saline and drug treatments are presented in current figure. Post-hoc results can be found in Table S5. Top row: cortical wide units. Middle row: cortical narrow units. Bottom row: pallidal units. Shadow indicates standard error of the mean. Legend includes count of all, oscillatory, and non-oscillatory sites, respectively.

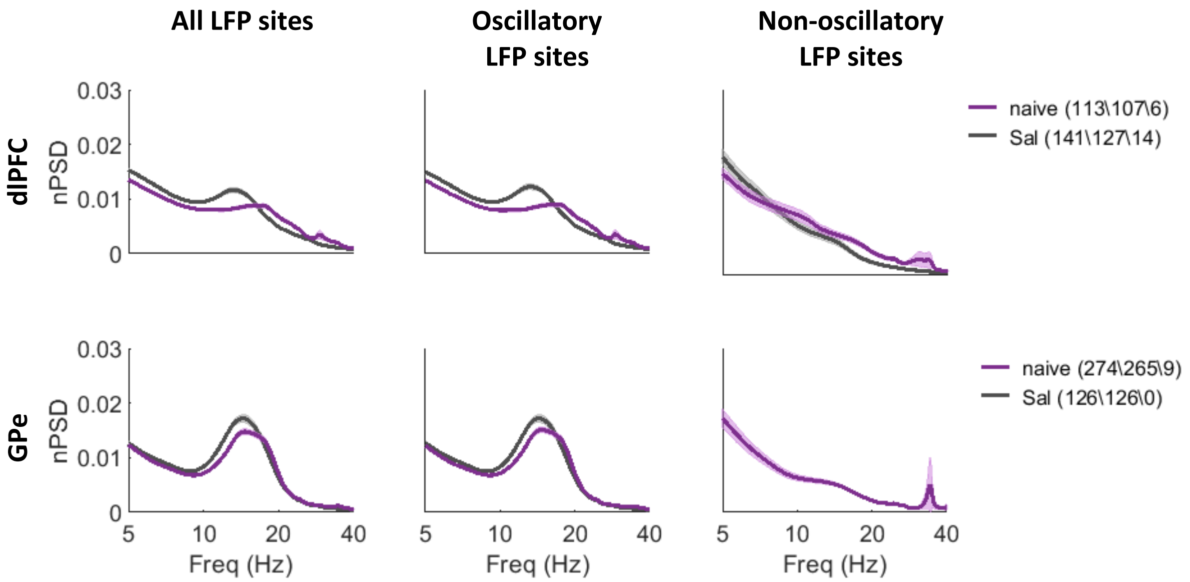


**Fig S4. Average nPSD of all (left), oscillatory (middle), and non-oscillatory (right) LFP sites in naïve and saline conditions.** Top row: dlPFC. Bottom row: GPe. Shadow indicates standard error of the mean. Legend indicates number of LFP sites in each category.

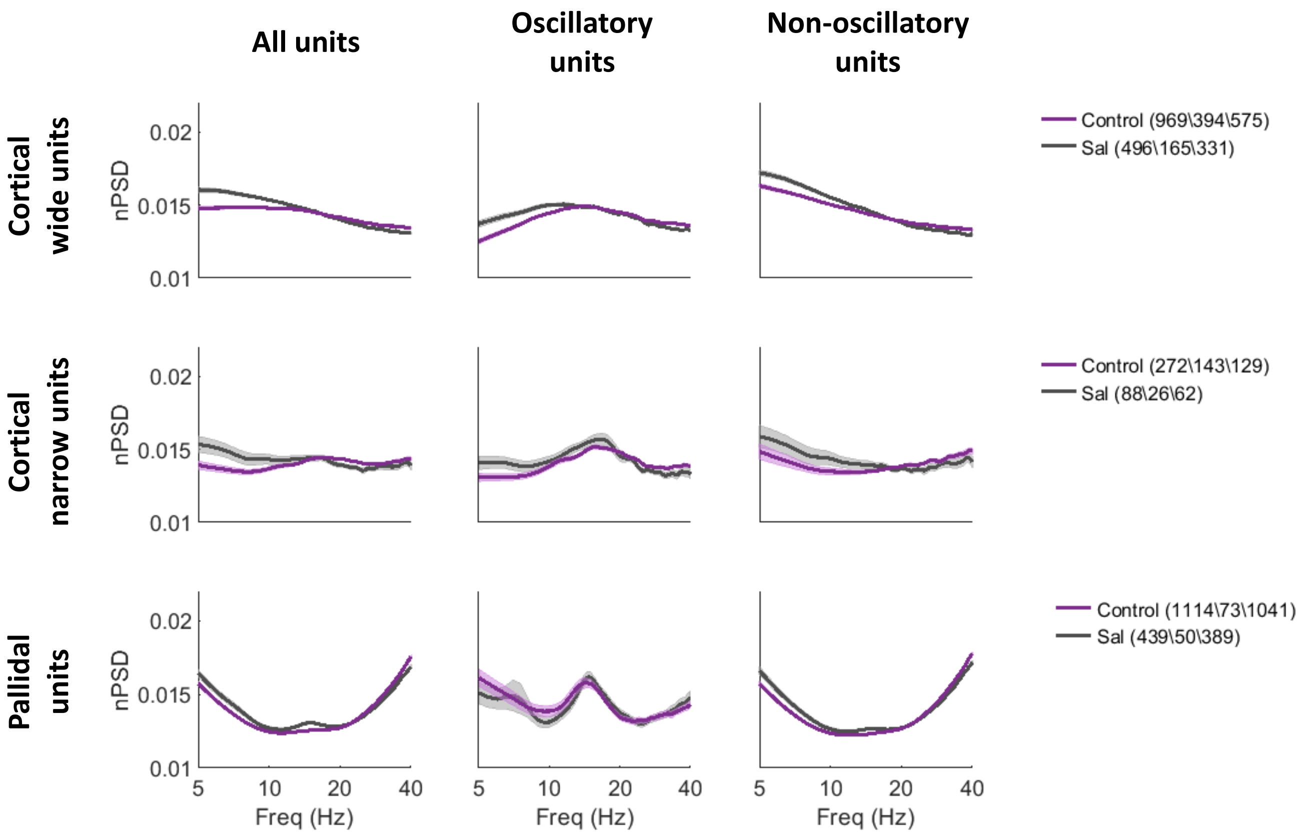


**Fig S5. Average nPSD of all (left), oscillatory (middle), and non-oscillatory (right) single units in naïve and saline conditions.** Top row: cortical wide units. Middle row: cortical narrow units. Bottom row: pallidal units. Shadow indicates standard error of the mean. Legend indicates number of units in each category.

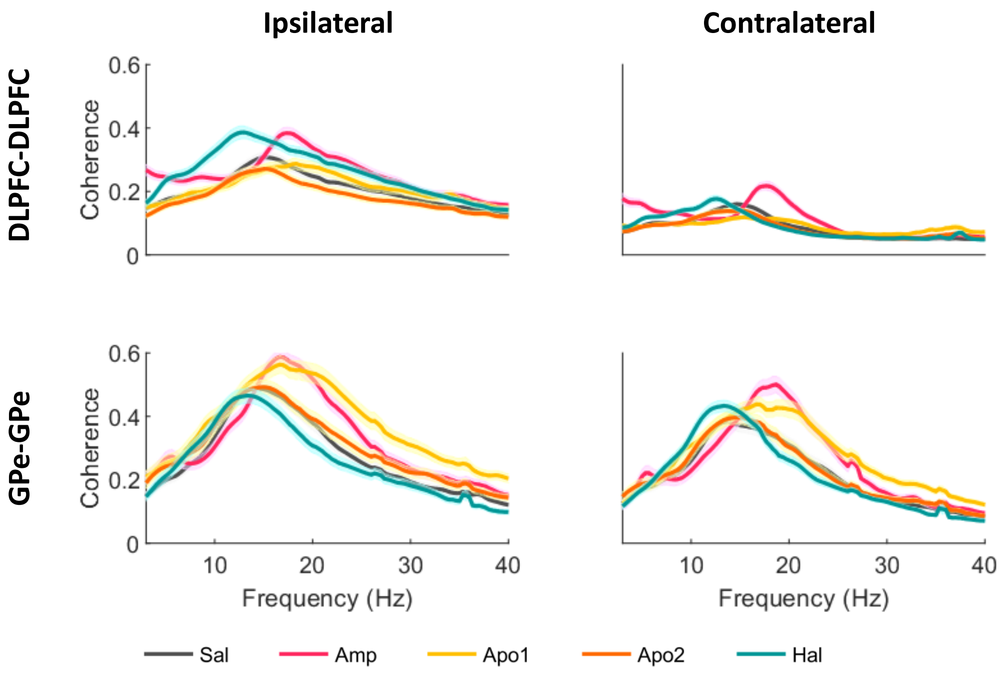


**Fig. S6: Coherence in the ipsilateral pairs is greater than in the contralateral pairs in the dlPFC, but not in the GPe.** Shadow indicates standard error of the mean.

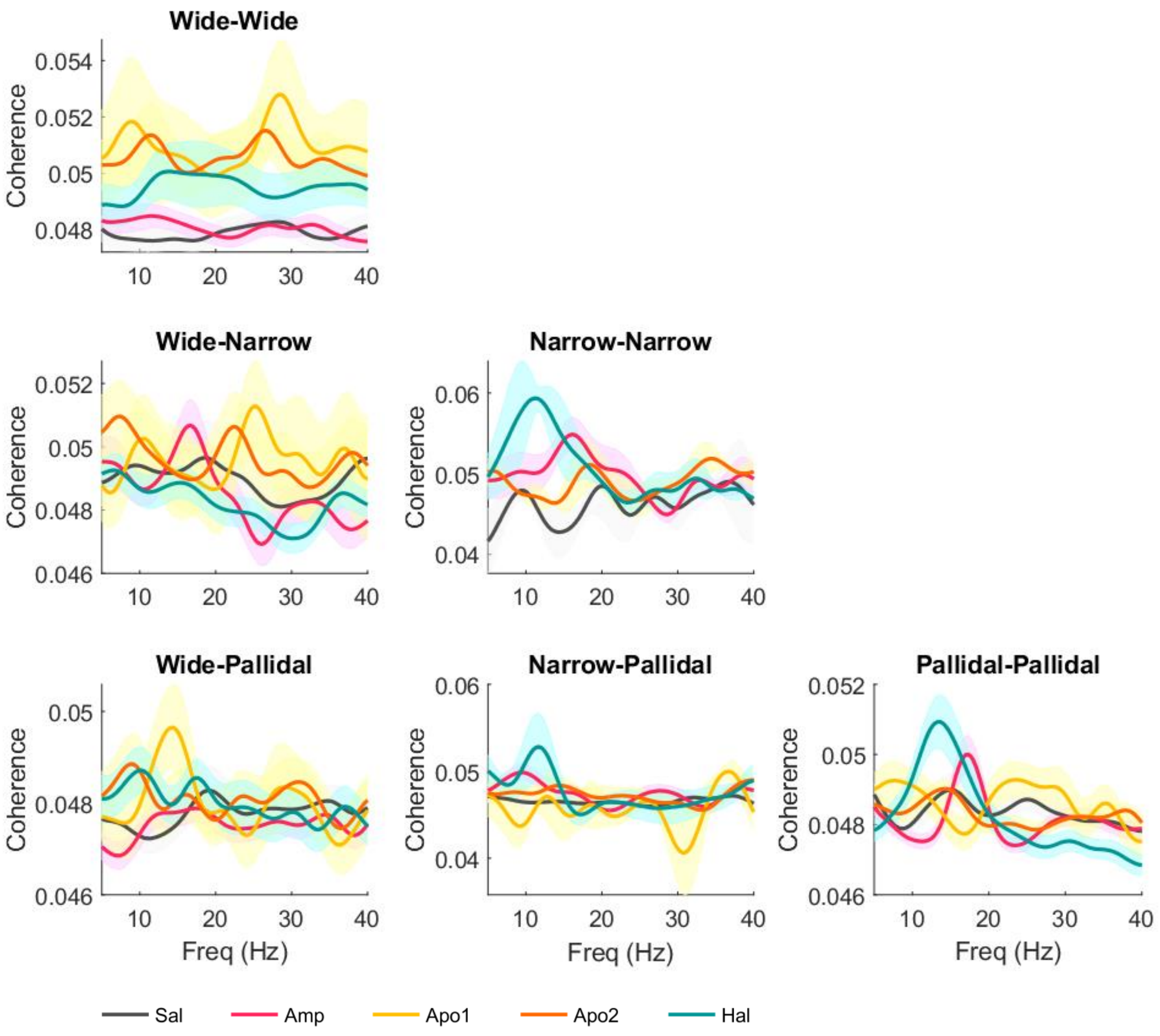


**Fig. S7: Single unit coherence in cortical narrow and pallidal pairs shows dopamine tone dependent shifts in beta frequency.** Shadow indicates standard error of the mean. Only unit pairs that were simultaneously recorded for at least five minutes were included in this analysis.

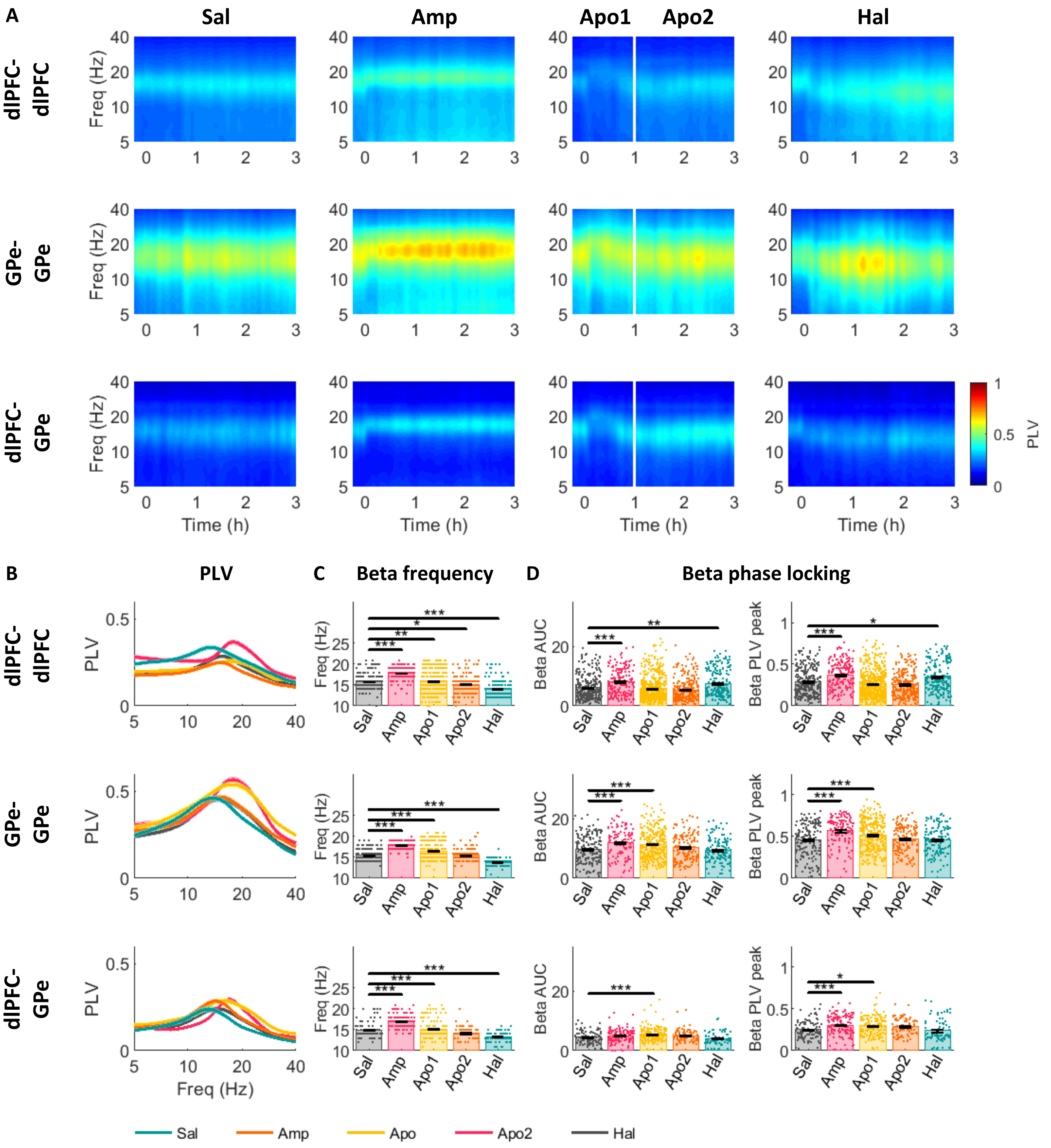


**Fig S8**. **Acute up- and down-modulation of dopamine tone up- and down-shifts the beta frequency of maximal PLV in LFP pairs within the CBG network of NHP.** (A) Average PLV of dlPFC-dlPFC (top), GPe-GPe (middle) and dlPFC-GPe (bottom) LFP pairs. Time 0 on x-axis indicates injection time. White line in the third column divides the post-apomorphine period into Apo1 -

agonistic phase and Apo2 – post-agonistic phase. (B-D) Properties of PLV beta peak in dlPFC-dlPFC (top), GPe-GPe (middle) and dlPFC-GPe (bottom) LFP pairs under each drug condition. (B) Average PLV (C) Frequency of PLV peaks in synchronized LFP sites (see methods) (D) Overall beta phase locking in the beta range was evaluated as area under the PLV curve (AUC) in 8-24Hz range (left) and as PLV peak within 8-24Hz frequency band (right). Bars indicate average values. Single points indicate individual unit value. Black vertical lines indicate standard error of the mean. Drug influence was evaluated by Kruskal-Wallis test followed by post-hoc pairwise comparisons. Comparisons between saline and drug treatments are presented in current figure. Full post-hoc results can be found in Table S7. * $p<0.05$ ** $p<0.01$ *** $p<0.001$

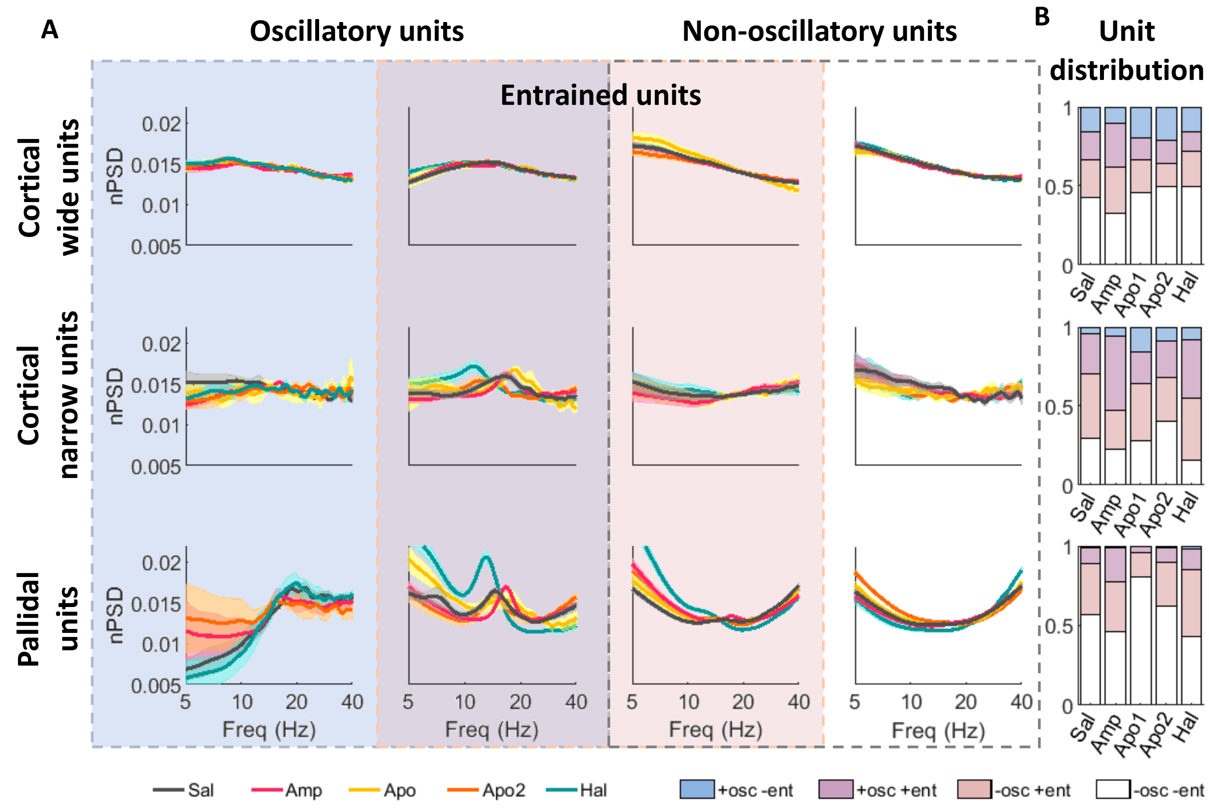


**Fig S9. Spectral activity of single units grouped by their oscillatory and LFP entrainment classification.** Units were classified in two independent processes as oscillatory or not, and as entrained to LFP beta activity or not (see methods). (A) Average nPSD of oscillatory and not entrained units (first column), oscillatory and entrained units (second column), non-oscillatory and entrained units (third column), non-oscillatory and not entrained units (fourth column). (B) Distribution of units into the aforementioned groups. Note the higher sensitivity of the entrainment analysis relative to the oscillation analysis. Entrainment analysis was based on LFP phase during spike occurrence, while oscillation analysis was based on beta peak prominence in the power spectrum. Top row: cortical wide units. Middle row: cortical narrow units. Bottom row: pallidal units. Shadow indicates standard error of the mean.

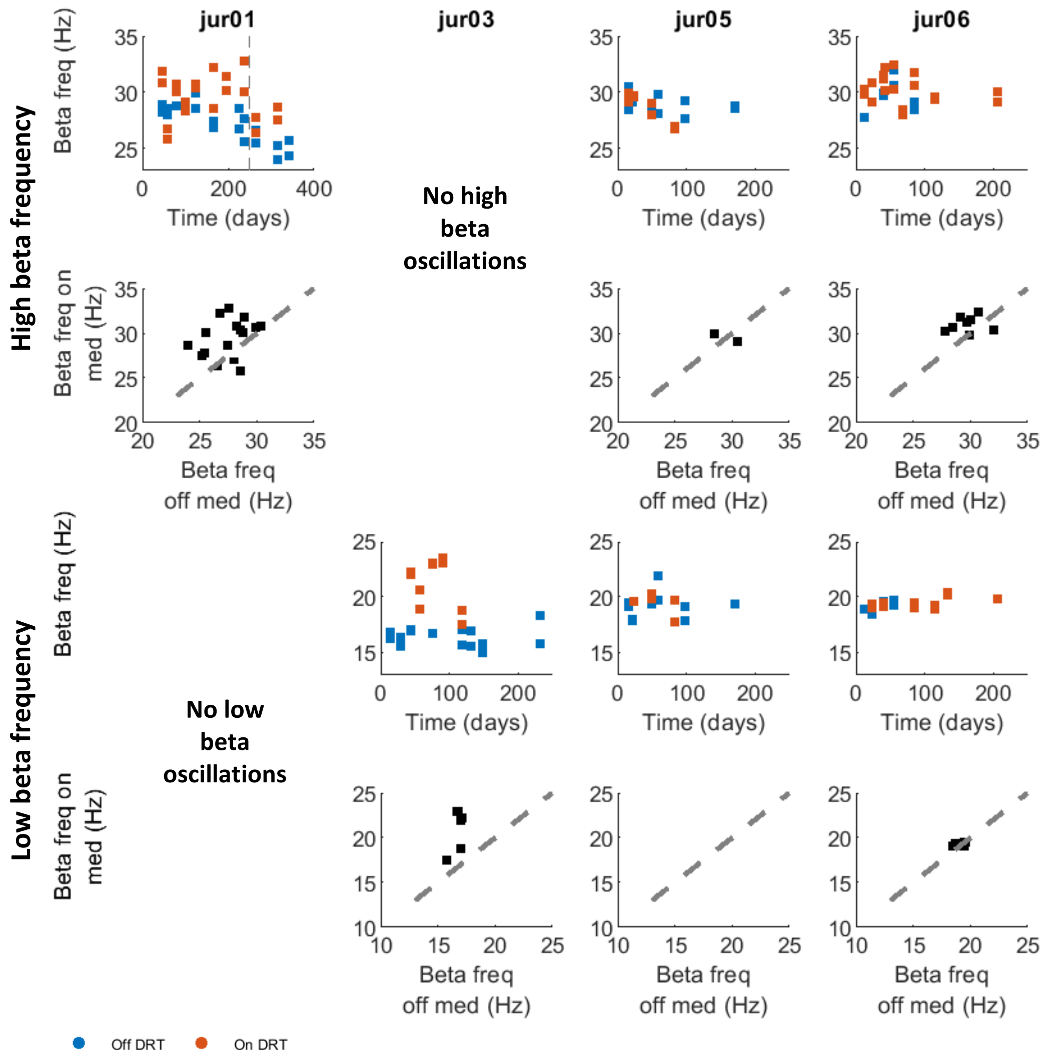


**Fig S10. Effect of acute and chronic dopamine modulation on LFP beta frequency in individual PD patients.** Each column shows data of a single patient. First row: Frequency of beta peak in the high beta domain as a function of time post-surgery. Each point represents average per day of beta peak frequency in one STN on (red) and off (blue) DRT. Second row: Comparison of beta frequency on and off DRT. Each point represents average per day of beta peak frequency in one STN in days with both off and on DRT sessions. X axis – off DRT. Y axis – on DRT. Clustering of data points above the diagonal line indicates a shift up in beta frequency in the on DRT condition relative to the off DRT condition. Third row: same as first row for low beta domain. Fourth row: same as second row for low beta domain. Patients can exhibit a peak in one or both beta domains. Gray dashed line indicates day 250 post-surgery. Recordings after this day were not included in the model to avoid exaggerated influence of jur01 data on MLEM results.

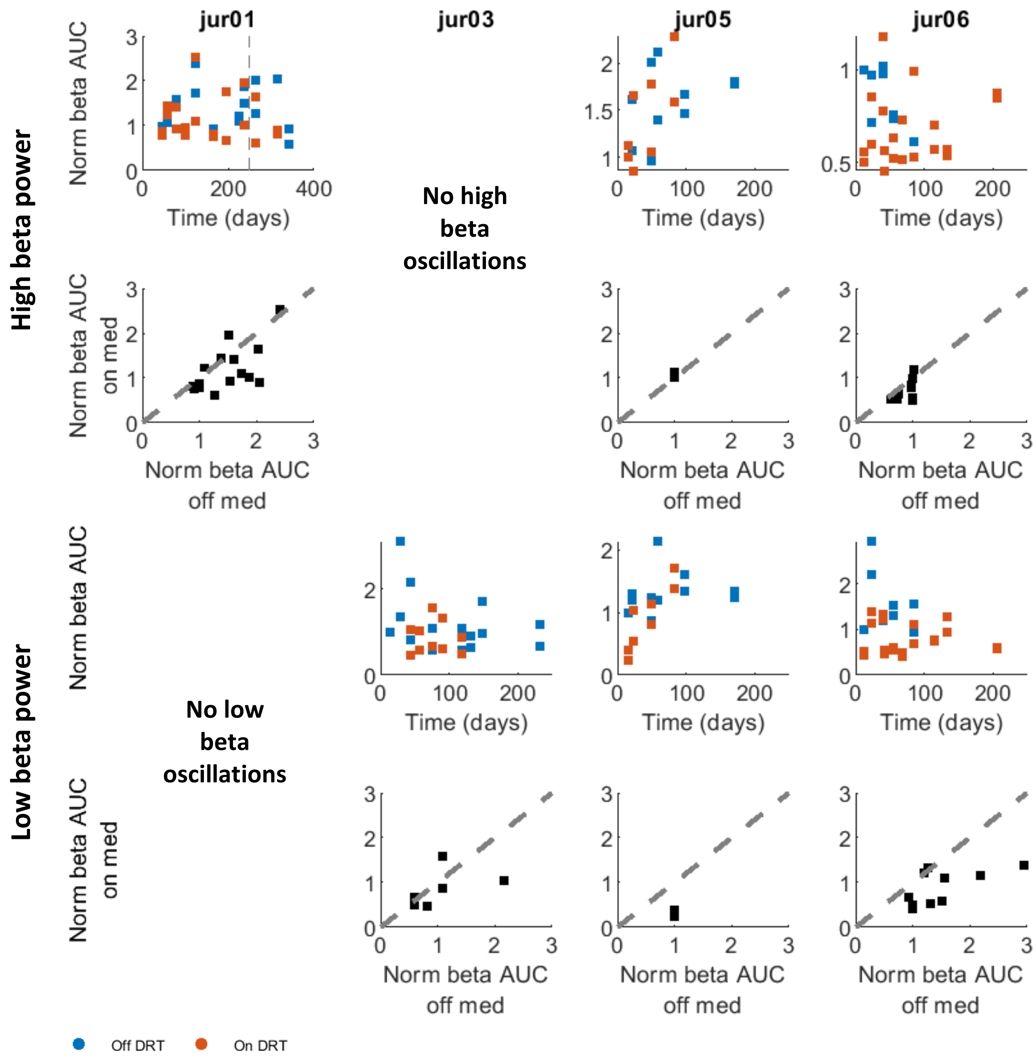


**Fig S11. Effect of acute and chronic dopamine modulation on LFP beta power in individual PD patients.** Beta power was evaluated as area under the curve (AUC) of the nPSD in the high and low beta bands. AUC values were normalized relative to beta AUC value in the first recording day after the surgery. Each column shows data of a single patient. First row: Beta power in the high-beta domain as a function of time post-surgery. Each point represents average per day of high-beta AUC on (red) and off (blue) DRT. Second row: Comparison of beta power on and off DRT. Each point represents average high-beta AUC in days with both off and on DRT sessions. X axis – off DRT. Y axis – on DRT. Clustering of data points below the diagonal line indicates a decrease in beta power in the on DRT condition relative to the off DRT condition. Third row: same as first row for low beta domain. Fourth row: same as second row for low beta domain. Patients can exhibit a peak in one or both beta domains. Gray dashed line indicates day 250 post-surgery. Recordings after this day were not included in the model to avoid exaggerated influence of jur01 data on MLEM results.

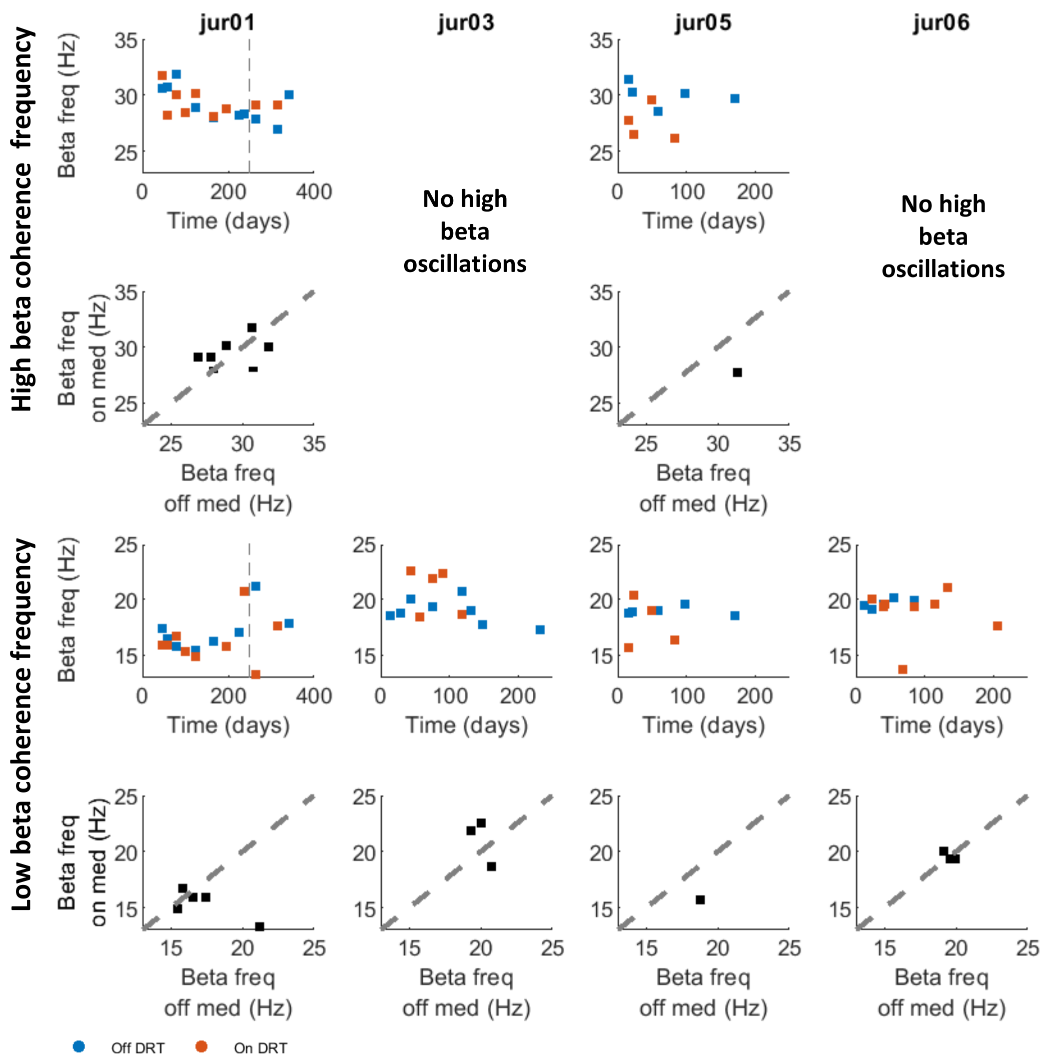


**Fig S12. Effect of acute and chronic dopamine modulation on LFP coherence beta frequency in individual PD patients.** Each column shows data of a single patient. First row: Frequency of beta coherence peak in the high beta domain as a function of time post-surgery. Each point represents average per day of beta coherence peak frequency on (red) and off (blue) DRT. Second row: Comparison of the frequency of beta coherence peak on and off DRT. Each dot represents average of beta coherence peak frequency in the high beta domain in days with both off and on DRT sessions. X axis – off DRT. Y axis – on DRT. Clustering of data points above the diagonal line indicates a shift up in beta coherence frequency in the on DRT condition relative to the off DRT condition. Third row: same as first row for low beta domain. Fourth row: same as second row for low beta domain. Patients can exhibit a peak in one or both beta domains. Gray dashed line indicates day 250 post-surgery. Recordings after this day were not included in the model to avoid exaggerated influence of jur01 data on MLEM results.

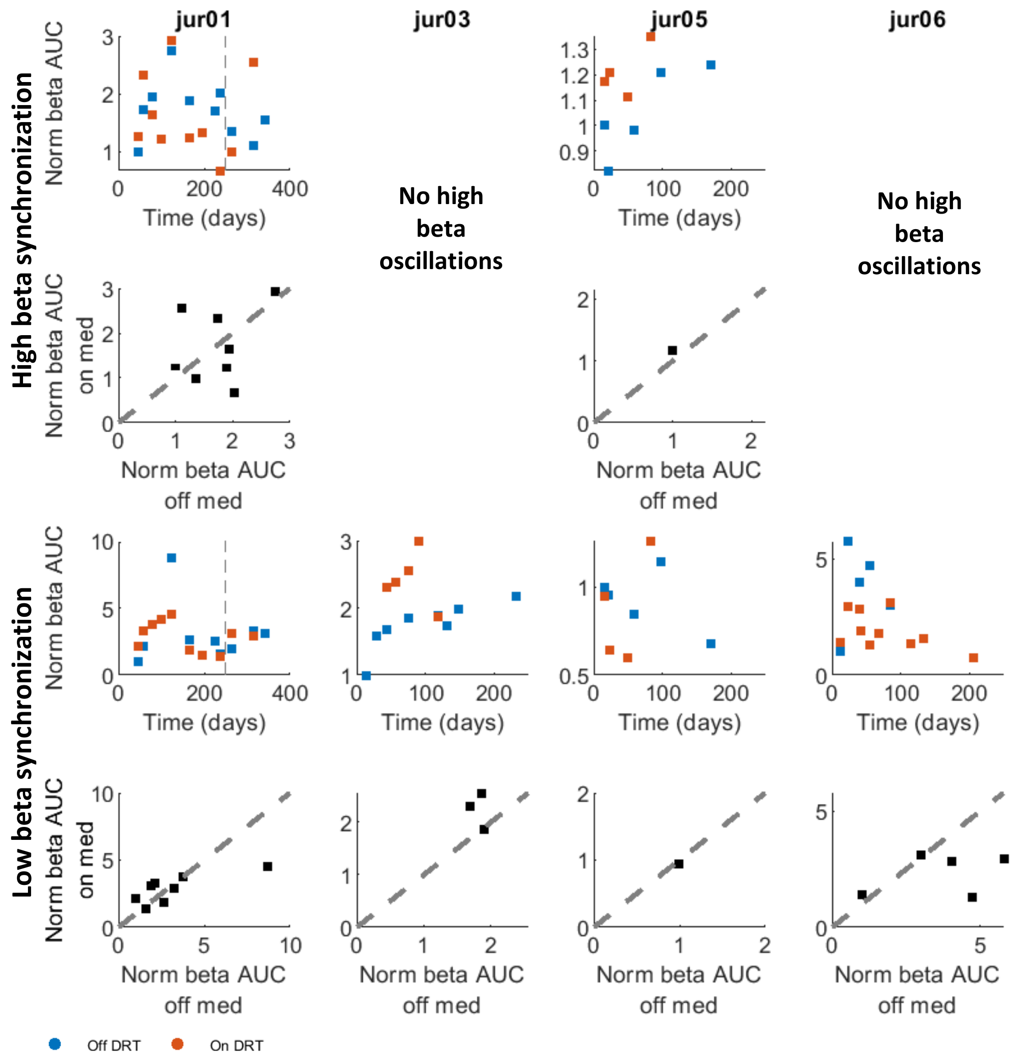


**Fig S13. Effect of acute and chronic dopamine modulation on beta synchrony in individual PD patients.** Beta synchrony is evaluated as area under the curve (AUC) of the coherence in the high and low beta domains. AUC values were normalized relative to beta AUC value in the first recording day after the surgery. Each column shows data of a single patient. First row: Beta synchrony in the high beta domain as a function of time post-surgery. Each point represents average per day of normalized beta AUC on (red) and off (blue) DRT. Second row: Comparison of beta synchrony on and off DRT. Each dot represents average per day of normalized beta AUC in the high-beta domain in days with both off and on DRT sessions. X axis – off DRT. Y axis – on DRT. Clustering of data points below the diagonal line indicates a decrease in beta synchrony in the on DRT condition relative to the off DRT condition. Third row: same as first row for low beta domain. Fourth row: same as second row for low beta domain. Patients can exhibit a peak in one or both beta domains. Gray dashed line indicates day 250 post-surgery. Recordings after this day were not included in the model to avoid exaggerated influence of jur01 data on MLEM results.

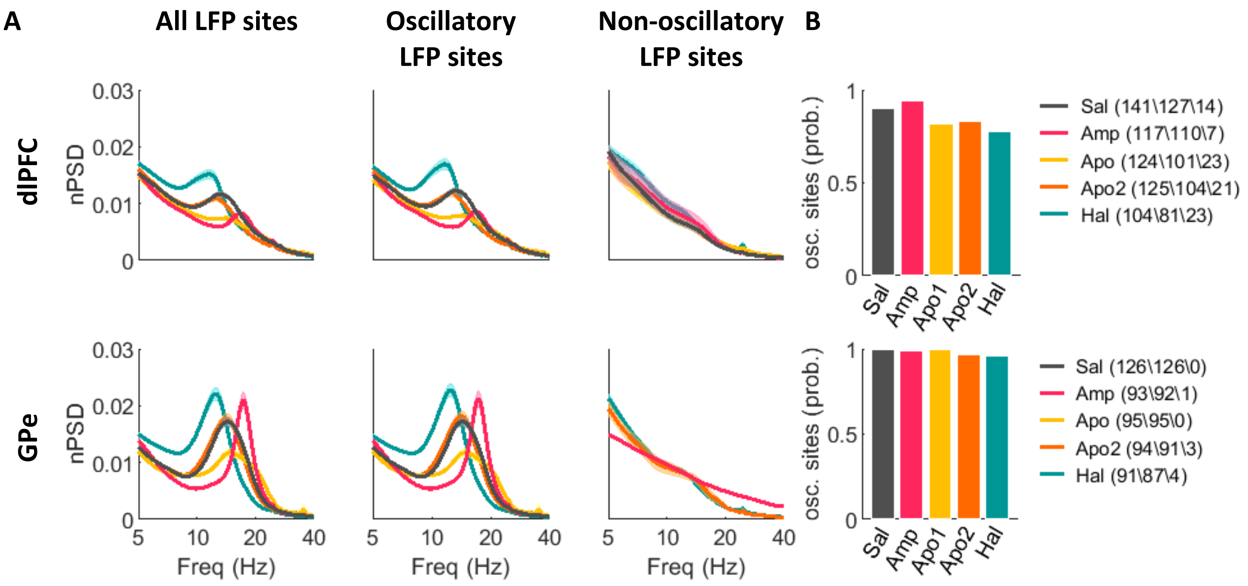


**Fig S14. Identification of LFP oscillatory sites.** (A) Average nPSD of all (left), oscillatory (middle), and non-oscillatory (right) LFP sites in all drug conditions. (B) Probability of oscillatory sites out of all the recorded sites in each condition. Drug effect was tested with chi square test (dlPFC: $\chi^2_{(4)}$=16.35, p=0.0026, GPe: $\chi^2_{(4)}$=9.77, p=0.0446) followed by pairwise comparisons with Bonferroni correction for multiple comparisons. No drug induced a significant change in oscillation probability relative to control (saline). Post-hoc results can be found in Table S5. Top row: dlPFC. Bottom row: GPe. Shadow indicates standard error of the mean. Legend includes count of all, oscillatory, and non-oscillatory sites, respectively.

### Supplementary Tables

| | | | **Sal** | **Amp** | **Apo (phase 1)** | **Apo (phase 2)** | **Hal** |
|---|---|---|---|---|---|---|---|
| **LFP sites** | Monkey G | dlPFC | 54 | 59 | 46 | 47 | 53 |
| | | GPe | 68 | 52 | 55 | 54 | 58 |
| | Monkey D | dlPFC | 87 | 58 | 78 | 78 | 51 |
| | | GPe | 58 | 41 | 40 | 40 | 53 |
| | **Total** | **dlPFC** | **141** | **117** | **124** | **125** | **104** |
| | | **GPe** | **126** | **93** | **95** | **94** | **91** |
| **Single units** | Monkey G | wide | 198 | 221 | 61 | 153 | 221 |
| | | narrow | 27 | 27 | 8 | 19 | 30 |
| | | HFD | 267 | 253 | 72 | 185 | 175 |
| | Monkey D | wide | 298 | 154 | 67 | 151 | 191 |
| | | narrow | 61 | 47 | 17 | 50 | 32 |
| | | HFD | 172 | 130 | 50 | 137 | 186 |
| | **Total** | **wide** | **496** | **375** | **128** | **304** | **412** |
| | | **narrow** | **88** | **74** | **25** | **69** | **62** |
| | | **HFD** | **439** | **383** | **122** | **322** | **361** |

**Table S1. Number of LFP sites and single units in dataset**

| Pts. | Number of recording days OFF/ON (both) DRT | Number of recording sessions OFF/ON DRT | Number of observations (sessions*sites) OFF/ON DRT |
|---|---|---|---|
| **Jur 01** | 10/11 (8) | 13/19 | 156/228 |
| **Jur 03** | 8/5 (3) | 10/6 | 120/72 |
| **Jur 05** | 6/4 (1) | 9/4 | 81/36 |
| **Jur 06** | 5/10 (5) | 11/19 | 132/228 |
| **Total** | 29/30 (17) | 43/48 | 489/564 |

**Table S2. Patient recording dataset.** Each recording day could have either only on DRT recording sessions, only off DRT sessions, or both. In column two, days with both on and off sessions are counted both as off day and as an on day

**A**

| | | dlPFC | | | | | GPe | | | | |
|---|---|---|---|---|---|---|---|---|---|---|---|
| | | Sal | Amp | Apo1 | Apo2 | Hal | Sal | Amp | Apo1 | Apo2 | Hal |
| LFP beta freq | mean | 14.41 | 16.74 | 15.71 | 13.52 | 12.54 | 14.47 | 16.84 | 15.82 | 14.51 | 12.34 |
| | SD | 2.8 | 1.48 | 3.5 | 2.25 | 2.95 | 1.72 | 1.07 | 3.08 | 2.18 | 1.08 |
| LFP Beta AUC | mean | 0.185 | 0.157 | 0.16 | 0.168 | 0.192 | 0.203 | 0.188 | 0.188 | 0.2 | 0.209 |
| | SD | 0.046 | 0.038 | 0.04 | 0.046 | 0.047 | 0.055 | 0.055 | 0.059 | 0.063 | 0.05 |
| LFP beta peak | mean | 0.012 | 0.009 | 0.008 | 0.011 | 0.014 | 0.019 | 0.022 | 0.015 | 0.019 | 0.023 |
| | SD | 0.006 | 0.004 | 0.003 | 0.006 | 0.009 | 0.008 | 0.013 | 0.007 | 0.009 | 0.011 |
| LFP beta peak (top 20%) | mean | --- | | | | | 0.023 | 0.015 | 0.013 | 0.021 | 0.029 |
| | SD | | | | | | 0.006 | 0.002 | 0.002 | 0.005 | 0.005 |

**B**

| | | dlPFC | | | | | | GPe | | | | | | | |
|---|---|---|---|---|---|---|---|---|---|---|---|---|---|---|---|
| | | LFP beta freq. | | LFP beta AUC | | LFP beta peak | | LFP beta freq. | | LFP beta AUC | | LFP beta peak | | LFP beta peak (top 20%) | |
| | | **p** | **g** | **p** | **g** | **p** | **g** | **p** | **g** | p | g | p | g | p | g |
| Sal | Amp | **9.9e-9** | **-1.01** | **2.5e-6** | **0.67** | **0.003** | **0.56** | **9.9e-9** | **-1.60** | ---. | | 0.820 | -0.27 | **1.6e-4** | **-1.92** |
| Sal | Apo1 | **0.023** | **-0.41** | **2.8e-5** | **0.57** | **1.1e-8** | **0.85** | **0.003** | **-0.56** | | | **0.003** | **0.59** | **0.033** | **1.73** |
| Sal | Apo2 | 0.061 | 0.34 | **0.017** | **0.36** | 0.138 | 0.24 | 0.999 | -0.02 | | | 1.000 | 0.03 | 0.991 | 0.32 |
| Sal | Hal | **1.6e-8** | **0.65** | 0.875 | -0.16 | 0.982 | -0.29 | **9.9e-9** | **1.42** | | | 0.364 | -0.35 | **0.002** | **-1.45** |
| Amp | Apo1 | **8.5e-4** | **0.38** | 0.984 | -0.09 | 0.067 | 0.40 | **2.4e-4** | **0.44** | | | **1.3e-4** | **0.73** | **9.9e-9** | **3.99** |
| Amp | Apo2 | **9.9e-9** | **1.69** | 0.234 | -0.28 | 0.696 | -0.29 | **9.9e-9** | **1.35** | | | 0.803 | 0.28 | **1.8e-5** | **3.09** |
| Amp | Hal | **9.9e-9** | **1.88** | **1.3e-7** | **-0.85** | **9.9e-4** | **-0.74** | **9.9e-9** | **4.17** | | | 0.957 | -0.05 | 0.977 | 0.27 |
| Apo1 | Apo2 | **6.9e-7** | **0.74** | 0.524 | -0.19 | **6.7e-4** | **-0.58** | **0.004** | **0.49** | | | **0.010** | **-0.52** | **0.113** | **-2.00** |
| Apo1 | Hal | **9.9e-9** | **0.97** | **1.4e-6** | **-0.75** | **1.1e-8** | **-0.96** | **9.9e-9** | **1.48** | | | **5.0e-6** | **-0.86** | **1.0e-8** | **-3.21** |
| apo2 | Hal | **0.003** | **0.38** | **0.001** | **-0.52** | 0.056 | -0.48 | **1.0e-8** | **1.25** | | | 0.373 | -0.36 | **2.8e-4** | **-2.16** |

**Table S3. Properties of LFP beta oscillations.** (A) Descriptive statistic (B) Post-hoc comparison results. p - p value, result of Tukey post-hoc test. g – effect size estimated by hedge's g. Comparisons that did not reach statistical significance and didn't require post-hoc test are marked with ---. Results are presented in Fig. 2

**A**

| | | Cortical wide units | | | | | Cortical narrow units | | | | | Pallidal units | | | | |
|---|---|---|---|---|---|---|---|---|---|---|---|---|---|---|---|---|
| | | sal | amp | apo1 | apo2 | hal | sal | amp | apo1 | apo2 | hal | sal | amp | apo1 | apo2 | hal |
| SUA beta freq. | Mean | 14.18 | 14.71 | 15.36 | 14.38 | 13.71 | 16.73 | 15.82 | 18.17 | 14.2 | 12.7 | 15.73 | 16.92 | 17.8 | 14.84 | 13.48 |
| | SD | 4.27 | 4.50 | 4.71 | 4.60 | 4.30 | 4.45 | 3.69 | 3.19 | 3.5 | 3.13 | 2.53 | 1.46 | 3.82 | 1.82 | 1.41 |
| SUA beta AUC | Mean | 0.078 | 0.078 | 0.081 | 0.08 | 0.08 | 0.071 | 0.071 | 0.07 | 0.066 | 0.072 | 0.043 | 0.043 | 0.033 | 0.04 | 0.045 |
| | SD | 0.016 | 0.012 | 0.016 | 0.015 | 0.016 | 0.017 | 0.006 | 0.014 | 0.011 | 0.017 | 0.013 | 0.013 | 0.01 | 0.012 | 0.017 |
| SUA beta peak | Mean | 0.0028 | 0.0028 | 0.003 | 0.0029 | 0.0029 | 0.0026 | 0.0026 | 0.0026 | 0.0024 | 0.0028 | 0.0014 | 0.0015 | 0.0011 | 0.0013 | 0.0016 |
| | SD | 0.0007 | 0.0005 | 0.0008 | 0.0007 | 0.0008 | 0.0009 | 0.0005 | 0.0006 | 0.0005 | 0.0009 | 0.0006 | 0.0006 | 0.0004 | 0.0005 | 0.0009 |

**B**

| | | Cortical wide units | | | | | | Cortical narrow units | | | | | | Pallidal units | | | | | |
|---|---|---|---|---|---|---|---|---|---|---|---|---|---|---|---|---|---|---|---|
| | | SUA beta freq. | | SUA beta AUC | | SUA beta peak | | SUA beta freq. | | SUA beta AUC | | SUA beta peak | | SUA beta freq. | | SUA beta AUC | | SUA beta peak | |
| | | p | g | p | g | p | g | p | g | p | g | p | g | **p** | **g** | p | g | p | g |
| Sal | Amp | | | | | 0.965 | 0.10 | 0.996 | -0.06 | 0.324 | 0.05 | | | **0.013** | **-0.17** | 0.997 | -0.01 | 0.527 | -0.08 |
| Sal | Apo1 | | | | | 0.338 | -0.20 | 0.338 | -0.44 | 1.000 | 0.06 | | | 0.224 | -0.20 | **9.9e-9** | **0.75** | **1.2e-8** | **0.61** |
| Sal | Apo2 | | | | | 0.119 | -0.13 | 0.965 | 0.12 | 0.361 | 0.35 | | | 0.998 | 0.01 | **0.017** | **0.25** | **0.040** | **0.22** |
| Sal | Hal | | | | | 0.996 | -0.05 | **0.033** | **0.48** | 0.993 | -0.02 | | | **6.3e-4** | **0.25** | 0.613 | -0.17 | 0.994 | -0.20 |
| Amp | Apo1 | --- | | --- | | 0.163 | -0.34 | 0.514 | -0.39 | 0.758 | 0.04 | --- | | 1.000 | -0.05 | **9.9e-9** | **0.73** | **9.9e-9** | **0.67** |
| Amp | Apo2 | | | | | **0.037** | **-0.26** | 0.853 | 0.18 | **0.004** | **0.50** | | | **0.011** | **0.18** | **0.009** | **0.25** | **3.0e-4** | **0.29** |
| Amp | Hal | | | | | 0.860 | -0.14 | **0.014** | **0.55** | 0.685 | -0.09 | | | **1.0e-8** | **0.41** | 0.825 | -0.16 | 0.813 | -0.13 |
| Apo1 | Apo2 | | | | | 1.000 | 0.08 | 0.153 | 0.56 | 0.659 | 0.34 | | | 0.173 | 0.20 | **2.4e-5** | **-0.54** | **3.2e-4** | **-0.44** |
| Apo1 | Hal | | | | | 0.514 | 0.15 | **0.001** | **0.92** | 1.000 | -0.09 | | | **1.9e-5** | **0.41** | **9.9e-9** | **-0.77** | **1.1e-8** | **-0.60** |
| Apo2 | Hal | | | | | 0.285 | 0.08 | 0.208 | 0.36 | 0.226 | -0.40 | | | **0.006** | **0.23** | **1.9e-4** | **-0.38** | **0.018** | **-0.36** |

**Table S4. Properties of SUA beta oscillations.** (A) Descriptive statistic (B) Post-hoc comparison results. p - p value, result of Tukey post-hoc test. g – effect size estimated by hedge's g. Comparisons that did not reach statistical significance and didn't require post-hoc test are marked with ---. Results are presented in Fig. 3

**A**

| | | dlPFC | GPe | Cortical wide units | Cortical narrow units | Pallidal units |
|---|---|---|---|---|---|---|
| Oscillatory site/unit probability | sal | 0.90 | 1.00 | 0.35 | 0.30 | 0.11 |
| | amp | 0.94 | 0.99 | 0.38 | 0.53 | 0.22 |
| | apo1 | 0.81 | 1.00 | 0.34 | 0.36 | 0.04 |
| | apo2 | 0.83 | 0.97 | 0.36 | 0.32 | 0.10 |
| | hal | 0.78 | 0.96 | 0.28 | 0.45 | 0.15 |

**B**

| | | dlPFC | | GPe | | Cortical wide units | | Cortical narrow units | | Pallidal units | |
|---|---|---|---|---|---|---|---|---|---|---|---|
| | | p | Φ | p | Φ | p | Φ | p | Φ | p | Φ |
| sal | amp | 1.0000 | -0.07 | 1.0000 | 0.08 | --- | | **0.0274** | **-0.24** | **3.0e-4** | **-0.15** |
| sal | apo1 | 0.4339 | -0.12 | -- | -- | | | 1.0000 | 0.06 | 0.1659 | -0.10 |
| sal | apo2 | 0.9802 | -0.10 | 0.4347 | -0.14 | | | 1.0000 | 0.03 | 1.0000 | -0.02 |
| sal | hal | 0.0846 | 0.17 | 0.1753 | -0.16 | | | 0.4975 | -0.16 | 1.0000 | 0.06 |
| amp | apo1 | **0.0315** | **-0.19** | 1.0000 | 0.07 | | | 1.0000 | -0.15 | **1.0e-4** | **-0.20** |
| amp | apo2 | 0.0857 | -0.17 | 1.0000 | -0.07 | | | 0.1189 | -0.21 | **1.0e-4** | **-0.16** |
| amp | hal | **0.0047** | **-0.24** | 1.0000 | -0.10 | | | 1.0000 | -0.08 | 0.1523 | -0.09 |
| apo1 | apo2 | 1.0000 | 0.02 | 0.7922 | -0.13 | | | 1.0000 | 0.04 | 0.4688 | -0.09 |
| apo1 | hal | 1.0000 | 0.04 | 0.3885 | -0.15 | | | 1.0000 | -0.08 | **0.0126** | **0.15** |
| apo2 | hal | 1.0000 | 0.07 | 1.0000 | -0.03 | | | 1.0000 | -0.14 | 0.3818 | 0.08 |

**Table S5. Probability of oscillatory sites/units.** Results are presented in Fig. S13 and Fig. S14

**A**

| | | dlPFC-dlPFC | | | | | GPe-GPe | | | | | dlPFC-GPe | | | | |
|---|---|---|---|---|---|---|---|---|---|---|---|---|---|---|---|---|
| | | Sal | Amp | Apo1 | Apo2 | Hal | Sal | Amp | Apo1 | Apo2 | Hal | Sal | Amp | Apo1 | Apo2 | Hal |
| LFP coherence beta frequency | mean | 15.39 | 17.3 | 15.89 | 15.16 | 13.74 | 14.96 | 17.53 | 16.48 | 15.23 | 13.19 | 14.28 | 16.69 | 14.77 | 13.49 | 13.09 |
| | SD | 2.07 | 1.73 | 3 | 2.55 | 2.09 | 1.73 | 1.29 | 2.56 | 2 | 0.62 | 2.12 | 1.38 | 2.53 | 1.18 | 1.8 |
| LFP coherence beta AUC | mean | 4.648 | 6.028 | 4.401 | 4.182 | 5.804 | 8.3 | 9.993 | 9.79 | 8.807 | 7.988 | 3.055 | 3.023 | 3.708 | 3.526 | 2.841 |
| | SD | 3.755 | 4.023 | 3.538 | 3.275 | 4.267 | 4.677 | 4.722 | 5.306 | 4.746 | 3.973 | 1.395 | 1.347 | 1.894 | 1.66 | 1.456 |
| LFP coherence beta peak | mean | 0.247 | 0.333 | 0.22 | 0.217 | 0.302 | 0.453 | 0.563 | 0.504 | 0.461 | 0.453 | 0.209 | 0.261 | 0.25 | 0.256 | 0.204 |
| | SD | 0.17 | 0.177 | 0.161 | 0.152 | 0.191 | 0.206 | 0.184 | 0.215 | 0.19 | 0.175 | 0.108 | 0.12 | 0.117 | 0.111 | 0.144 |

**B**

| | | dlPFC-dlPFC | | | | | | Gpe-GPe | | | | | | dlPFC-GPe | | | | | |
|---|---|---|---|---|---|---|---|---|---|---|---|---|---|---|---|---|---|---|---|
| | | beta freq. | | beta AUC | | Beta peak | | beta freq. | | beta AUC | | Beta peak | | beta freq. | | beta AUC | | Beta peak | |
| | | p | g | p | g | p | g | p | g | p | g | p | g | p | g | p | g | p | g |
| Sal | Amp | **9.9e-9** | **-0.98** | **5.6e-5** | **-0.36** | **6.0e-7** | **-0.50** | **9.9e-9** | **-1.64** | **0.033** | **-0.36** | **4.0e-5** | **-0.56** | **9.9e-9** | **-1.38** | 1.000 | 0.02 | **0.004** | **-0.45** |
| Sal | Apo1 | **5.8e-5** | **-046** | 0.975 | 0.01 | 0.405 | 0.14 | **9.9e-9** | **-1.31** | **0.001** | **-0.47** | **0.001** | **-0.43** | **2.9e-5** | **-0.74** | **0.003** | **-0.48** | 0.258 | -0.31 |
| Sal | Apo2 | 0.711 | 0.10 | 0.779 | 0.13 | 0.324 | 0.18 | 0.851 | -0.14 | 0.964 | -0.11 | 1.000 | -0.04 | 0.160 | 0.44 | 0.260 | -0.31 | **0.040** | **-0.44** |
| Sal | Hal | **9.9e-9** | **0.79** | **0.004** | **-0.29** | **0.007** | **-0.31** | **9.9e-9** | **1.30** | 0.980 | 0.07 | 0.999 | -0.00 | **7.8e-4** | **0.59** | 0.780 | 0.15 | 0.791 | 0.04 |
| Amp | Apo1 | **1.2e-6** | **0.24** | **7.7e-4** | **0.36** | **1.0e-8** | **0.63** | 0.837 | -0.09 | 0.966 | -0.14 | 0.871 | 0.08 | **0.013** | **0.28** | **0.001** | **-0.52** | 0.807 | 0.14 |
| Amp | Apo2 | **9.9e-9** | **0.95** | **3.8e-7** | **0.51** | **9.9e-9** | **0.71** | **9.9e-9** | **1.34** | 0.197 | 0.25 | **1.7e-4** | **0.55** | **9.9e-9** | **2.44** | 0.184 | -0.34 | 0.998 | 0.04 |
| Amp | Hal | **9.9e-9** | **1.84** | 0.835 | 0.05 | 0.282 | 0.17 | **9.9e-9** | **4.40** | **0.010** | **0.46** | **3.5e-5** | **0.62** | **9.9e-9** | **2.32** | 0.773 | 0.13 | **1.2e-4** | **0.44** |
| Apo1 | Apo2 | **1.3e-7** | **0.51** | 0.420 | 0.12 | 1.000 | 0.04 | **9.9e-9** | **1.10** | **0.024** | **0.37** | **0.004** | **0.40** | **1.0e-8** | **1.20** | 0.541 | 0.20 | 0.952 | -0.10 |
| Apo1 | Hal | **9.9e-9** | **1.03** | **0.031** | **-0.30** | **9.3e-6** | **-0.44** | **9.9e-9** | **2.48** | **3.5e-4** | **0.56** | **8.8e-4** | **0.45** | **9.9e-9** | **1.25** | **9.4e-5** | **0.58** | **0.027** | **0.30** |
| Apo2 | Hal | **1.1e-8** | **0.60** | **7.0e-5** | **-0.43** | **4.7e-6** | **-0.50** | **9.9e-9** | **1.36** | 0.768 | 0.19 | 0.995 | 0.04 | 0.454 | 0.26 | **0.025** | **0.44** | **0.002** | **0.41** |

**Table S6. Properties of LFP beta coherence.** (A) Descriptive statistic (B) Post-hoc comparison results. p - p value, result of Tukey post-hoc test. g – effect size estimated by hedge's g. Results are presented in Fig. 4

**A**

| | | dlPFC-dlPFC | | | | | GPe-GPe | | | | | dlPFC-GPe | | | | |
|---|---|---|---|---|---|---|---|---|---|---|---|---|---|---|---|---|
| | | Sal | Amp | Apo1 | Apo2 | Hal | Sal | Amp | Apo1 | Apo2 | Hal | Sal | Amp | Apo1 | Apo2 | Hal |
| LFP coherence beta frequency | mean | 15.781 | 17.846 | 15.881 | 15.212 | 14.033 | 15.519 | 17.975 | 16.559 | 15.537 | 13.852 | 14.971 | 17.044 | 15.22 | 14.226 | 13.382 |
| | SD | 1.602 | 1.408 | 2.354 | 1.8 | 1.892 | 1.444 | 0.908 | 1.985 | 1.346 | 0.725 | 1.812 | 1.48 | 2.072 | 1.283 | 1.211 |
| LFP coherence beta AUC | mean | 6.115 | 8.06 | 5.808 | 5.397 | 7.426 | 9.73 | 11.996 | 11.445 | 10.253 | 9.526 | 4.487 | 4.959 | 5.446 | 5.015 | 4.1 |
| | SD | 4.084 | 4.278 | 4.143 | 3.737 | 4.551 | 4.735 | 4.512 | 4.924 | 4.354 | 3.931 | 1.98 | 2.093 | 2.364 | 2.014 | 2.051 |
| LFP coherence beta peak | mean | 0.29 | 0.374 | 0.264 | 0.254 | 0.343 | 0.461 | 0.57 | 0.514 | 0.47 | 0.46 | 0.25 | 0.306 | 0.296 | 0.289 | 0.242 |
| | SD | 0.159 | 0.161 | 0.161 | 0.147 | 0.173 | 0.193 | 0.165 | 0.18 | 0.159 | 0.16 | 0.106 | 0.118 | 0.109 | 0.097 | 0.136 |

**B**

| | | dlPFC-dlPFC | | | | | | Gpe-GPe | | | | | | dlPFC-GPe | | | | | |
|---|---|---|---|---|---|---|---|---|---|---|---|---|---|---|---|---|---|---|---|
| | | beta freq. | | beta AUC | | Beta peak | | beta freq. | | beta AUC | | Beta peak | | beta freq. | | beta AUC | | Beta peak | |
| | | p | g | p | g | p | g | p | g | p | g | p | g | p | g | p | g | p | g |
| Sal | Amp | **9.9e-9** | **-1.35** | **5.1e-6** | **-0.47** | **2.1e-6** | **-0.52** | **9.9e-9** | **-1.95** | **7.8e-4** | **-0.48** | **5.3e-6** | **-0.60** | **9.9e-9** | **-1.27** | 0.339 | -0.23 | **7.6e-4** | **-0.49** |
| Sal | Apo1 | **0.007** | **-0.36** | 1.000 | -0.03 | 0.512 | 0.10 | **9.9e-9** | **-1.17** | **1.9e-5** | **-0.58** | **2.4e-4** | **-0.51** | **2.9e-4** | **-0.64** | **1.7e-4** | **-0.62** | **0.014** | **-0.47** |
| Sal | Apo2 | **0.019** | **0.33** | 0.214 | 0.18 | 0.082 | 0.24 | 0.996 | -0.01 | 0.969 | -0.11 | 1.000 | -0.05 | 0.112 | 0.46 | 0.525 | -0.26 | 0.131 | -0.38 |
| Sal | Hal | **9.9e-9** | **1.01** | **0.010** | **-0.31** | **0.012** | **-0.32** | **9.9e-9** | **1.43** | 0.971 | 0.05 | 0.982 | 0.00 | **1.8e-6** | **1.00** | 0.609 | 0.19 | 0.942 | 0.07 |
| Amp | Apo1 | **1.0e-8** | **0.58** | **5.5e-6** | **0.42** | **1.1e-8** | **0.59** | 0.152 | 0.26 | 0.987 | -0.12 | 0.844 | 0.07 | **0.011** | **0.43** | **0.039** | **-0.41** | 0.998 | 0.02 |
| Amp | Apo2 | **9.9e-9** | **1.59** | **1.0e-8** | **0.67** | **9.9e-9** | **0.78** | **9.9e-9** | **2.08** | **0.012** | **0.39** | **8.0e-6** | **0.61** | **9.9e-9** | **1.99** | 1.000 | 0.03 | 0.737 | 0.15 |
| Amp | Hal | **9.9e-9** | **2.27** | 0.441 | 0.14 | 0.316 | 0.18 | **9.9e-9** | **5.08** | **1.6e-4** | **0.59** | **1.4e-6** | **0.67** | **9.9e-9** | **2.63** | **0.012** | **0.41** | **1.1e-4** | **0.51** |
| Apo1 | Apo2 | **1.4e-8** | **0.59** | 0.257 | 0.20 | 0.881 | 0.12 | **9.9e-9** | **1.17** | **8.4e-4** | **0.49** | **3.1e-4** | **0.50** | **1.4e-8** | **1.12** | **0.066** | **0.38** | 0.929 | 0.13 |
| Apo1 | Hal | **9.9e-9** | **1.07** | **0.009** | **-0.27** | **4.3e-5** | **-0.40** | **9.9e-9** | **2.43** | **3.5e-6** | **0.67** | **6.5e-5** | **0.56** | **9.9e-9** | **1/59** | **9.4e-7** | **0.76** | **0.002** | **0.47** |
| Apo2 | Hal | **7.8e-8** | **0.64** | **2.7e-6** | **-0.49** | **4.5e-7** | **0.56** | **9.9e-9** | **1.57** | 0.752 | 0.17 | 0.995 | 0.06 | **0.047** | **0.67** | **0.040** | **0.45** | **0.030** | **0.40** |

**Table S7. Properties of LFP beta PLV.** (A) Descriptive statistic (B) Post-hoc comparison results. p - p value, result of Tukey post-hoc test. g – effect size estimated by hedge's g. Results are presented in Fig. S

**A**

| | | | Wide | | | | | narrow | | | | | HFD | | | | |
|---|---|---|---|---|---|---|---|---|---|---|---|---|---|---|---|---|---|
| | | | Sal | Amp | Apo1 | Apo2 | Hal | Sal | Amp | Apo1 | Apo2 | Hal | Sal | Amp | Apo1 | Apo2 | Hal |
| Ent. Probability | | | 0.45 | 0.57 | 0.37 | 0.31 | 0.38 | 0.66 | 0.73 | 0.59 | 0.53 | 0.78 | 0.42 | 0.52 | 0.19 | 0.37 | 0.53 |
| Vector length | | mean | 0.1 | 0.095 | 0.095 | 0.085 | 0.087 | 0.101 | 0.122 | 0.079 | 0.083 | 0.111 | 0.018 | 0.021 | 0.008 | 0.015 | 0.023 |
| | | SD | 0.07 | 0.065 | 0.063 | 0.057 | 0.062 | 0.075 | 0.086 | 0.044 | 0.048 | 0.073 | 0.016 | 0.024 | 0.007 | 0.017 | 0.022 |
| Preferred phase | Group by drugs | mean | -2.83 | -2.82 | -2.96 | -2.83 | -2.82 | -2.11 | -2.1 | -2.47 | -2.04 | -2.2 | 2.24 | 3.13 | 1.43 | 1.09 | -0.42 |
| | | ang. SD | 0.65 | 0.63 | 0.6 | 0.86 | 0.87 | 1.06 | 0.74 | 0.64 | 0.85 | 1.27 | 1.37 | 1.17 | 1.03 | 1.34 | 1.35 |
| | | | Low beta | | High beta | | | Low beta | | High beta | | | Low beta | | High beta | | |
| | Group by beta | mean | -2.82 | | -2.82 | | | -1.66 | | -2.19 | | | 0.58 | | 2.68 | | |
| | | ang. SD | 0.62 | | 0.68 | | | 1.11 | | 1.04 | | | 1.31 | | 1.31 | | |
| | Group by beta | Mean | -2.81 | | -2.83 | | | -1.94 | | -2.2 | | | 0.77 | | 2.92 | | |
| | | ang. SD | 0.75 | | 0.71 | | | 1.16 | | 0.87 | | | 1.3 | | 1.3 | | |

**B**

| | | wide | | | | | narrow | | | | | HFD | | | | |
|---|---|---|---|---|---|---|---|---|---|---|---|---|---|---|---|---|
| | | Entrainment probability | | Vector length | | Pref. phase | Entrainment probability | | Vector length | | Pref. phase | Entrainment probability | | Vector length | | Pref. phase |
| | | p | Φ | p | g | p | p | Φ | p | g | p | p | Φ | p | g | p |
| Sal | Amp | **0.0049** | **-0.12** | 0.955 | 0.07 | | 1.0000 | -0.08 | | | | 0.0749 | -0.10 | 1.0000 | -0.15 | 0.274 |
| Sal | Apo1 | 1.0000 | -0.06 | 0.996 | 0.08 | | 1.0000 | -0.06 | | | | **1.9e-4** | **-0.20** | **9.9e-9** | **0.68** | 0.273 |
| Sal | Apo2 | **0.0017** | **-0.14** | 0.087 | 0.23 | | 1.0000 | -0.13 | | | | 1.0000 | -0.05 | **0.0047** | **0.16** | 0.359 |
| Sal | Hal | 0.4696 | 0.07 | 0.054 | 0.20 | | 1.0000 | -0.13 | | | | **0.0452** | **0.11** | **0.0305** | **-0.25** | 0.486 |
| Amp | Apo1 | **0.0018** | **-0.17** | 1.000 | 0.01 | --- | 1.0000 | -0.13 | --- | | --- | **2.5e-8** | **-0.29** | **9.9e-9** | **0.60** | 0.110 |
| Amp | Apo2 | **3.7e-10** | **-0.27** | 0.401 | 0.17 | | 0.1523 | -0.21 | | | | **0.0017** | **-0.15** | **0.0049** | **0.27** | 0.123 |
| Amp | Hal | **2.0e-6** | **-0.19** | 0.343 | 0.13 | | 1.0000 | 0.06 | | | | 1.0000 | 0.01 | **0.0442** | **-0.07** | 0.274 |
| Apo1 | Apo2 | 2.0971 | 0.06 | 0.640 | 0.17 | | 1.0000 | 0.05 | | | | **0.0071** | **-0.17** | **2.0e-5** | **-0.48** | 0.273 |
| Apo1 | Hal | 1.0000 | -0.01 | 0.623 | 0.12 | | 0.8143 | -0.19 | | | | **1.5e-8** | **0.31** | **9.9e-9** | **-0.76** | 0.597 |
| Apo2 | Hal | 0.5619 | -0.07 | 1.000 | -0.04 | | **0.0320** | **-0.26** | | | | **0.0010** | **0.16** | **2.5e-8** | **-0.37** | 0.538 |

**Table S8. Properties of SUA beta entrainment.** (A) Descriptive statistic (B) Post-hoc comparison results. p - p value, result of Tukey post-hoc test. g – effect size estimated by hedge's g. Φ – effect size estimated by phi coefficient. Comparisons that did not reach statistical significance and didn't require post-hoc test are marked with ---. Results are presented in Fig. 5

| Pts. | Age | Gender | Duration of disease (years) | Baseline Medications (dose) | Levodopa Equivalent Dose (LED) | Baseline UPDRS III motor score (PD) | DBS Lead Target [x,y,z] | Optimal stimulation parameters: (Frequency (Hz); Pulse Duration (µs); Contact configuration; Voltage(V)) |
|---|---|---|---|---|---|---|---|---|
| **Jur 01** | 66 | F | 8 | Stalevo 50 mg q4d<br>Rasagiline 2 mg qid | 1066 | 35 | Left:<br>[-12, -4, -4.5]<br>Right:<br>[12.25, -3.5, -5.5] | Left:<br>(180; 60; c+9-; 2)<br>Right:<br>(180; 60; c+1-; 1.9) |
| **Jur 03** | 54 | M | 9 | Carbidopa 12.5 mg q3h<br>Levodopa 125 mg q3h<br>Biperiden 1mg q4h<br>Ropinirole 4mg qid | 1257.5 | 31 | Left:<br>[-11.25, -1.75, -4.5]<br>Right:<br>[11, -2.5, -5] | Left:<br>(130; 60; c+9-; 2.1)<br>Right:<br>(130; 60; c+1-; 1.6) |
| **Jur 05** | 52 | F | 10 | Carbidopa 25 mg q6h Levodopa 250 mg q6h | 1125 | 42 | Left:<br>[-10.5, -2.5, -4]<br>Right:<br>[10.75, -3.5, -5] | Left:<br>(130; 60; c+8-11-; 1.9)<br>Right:<br>(130; 60; c+1-2-; 1.3) |
| **Jur 06** | 52 | F | 9 | Carbidopa 25 mg q5h Levodopa 250 mg q5h<br>Rasagiline 1mg qid<br>Ropinirole 8mg qid | 2165 | 43 | Left:<br>[-11, -3, -4]<br>Right:<br>[11.5, -3, -4.5] | Left:<br>(130; 60; c+9-11-; 2.2)<br>Right:<br>(130; 60; c+1-3-; 1.8) |

**Table S9: Patient demographics and treatment**

| | Dependent variable | Factor | Estimated coefficient | F | DF 1 | DF 2 | p |
|---|---|---|---|---|---|---|---|
| Beta frequency/ power | High –beta frequency | **Time** | **-0.0131** | **33.949** | **1** | **462** | **1.1e-08** |
| | | DRT | -0.4710 | 2.1604 | 1 | 462 | 0.1423 |
| | | **Time x DRT** | **0.0146[a]** | **25.36** | **1** | **462** | **6.8e-07** |
| | low-beta frequency | Time | 0.0013 | 0.101 | 1 | 414 | 0.7508 |
| | | DRT | 2.0924 | 3.736 | 1 | 414 | 0.0539 |
| | | Time x DRT | -0.0058[a] | 1.1366 | 1 | 414 | 0.287 |
| | High-beta AUC | Time | 0.0014 | 0.7346 | 1 | 736 | 0.3917 |
| | | DRT | -0.1868 | 3.3476 | 1 | 736 | 0.0677 |
| | | Time x DRT | 0.0014[a] | 3.558 | 1 | 736 | 0.0597 |
| | Low-beta AUC | Time | -0.0023 | 1.4921 | 1 | 665 | 0.2223 |
| | | **DRT** | **-0.6771** | **40.934** | **1** | **665** | **2.9e-10** |
| | | **Time x DRT** | **0.0039[a]** | **6.3891** | **1** | **665** | **0.0117** |
| Beta coherence | High –beta frequency | **Time** | **-0.0132** | **8.0614** | **1** | **120** | **0.0053** |
| | | DRT | -2.0999 | 3.266 | 1 | 120 | 0.0732 |
| | | Time x DRT | 0.0066[a] | 0.4526 | 1 | 120 | 0.5024 |
| | low-beta frequency | Time | -0.0024 | 0.5466 | 1 | 201 | 0.4606 |
| | | DRT | -0.5080 | 0.6182 | 1 | 201 | 0.4327 |
| | | Time x DRT | 0.0077[a] | 2.0826 | 1 | 201 | 0.1505 |
| | High-beta AUC | Time | 0.0022 | 1.9532 | 1 | 200 | 0.1638 |
| | | DRT | 0.4627 | 2.5503 | 1 | 200 | 0.1119 |
| | | Time x DRT | -0.0056[a] | 5.6426 | 1 | 200 | 0.0185 |
| | Low-beta AUC | Time | 0.0005 | 0.0221 | 1 | 476 | 0.8819 |
| | | DRT | 0.3424 | 0.2927 | 1 | 476 | 0.5887 |
| | | Time x DRT | -0.0090[a] | 3.7752 | 1 | 476 | 0.0526 |

**Table S10: Time and DRT effects on beta properties in PD patients.** To estimate the contribution of time and DRT (dopamine replacement therapy) on beta properties in PD patients, a mixed linear effect model (MLEM) was constructed for each dependent variable (column 1 and 2). The model included fixed effect terms for time, DRT and their interaction (column 3). The resulted estimated coefficients are presented in column 4. One-way ANOVA was used on model output to determine the significance of each factor. The test examines significance of the fixed factors beyond levels, which was important for the interaction factor. ANOVA results are presented in the columns 5-8. Note that a separate model was constructed for high-beta and low-beta properties. Subjects were clustered as having low-beta, high-beta or both and traces were included in the analysis accordingly. i.e. If a subject had low-beta, all his traces were included in the low-beta analysis and the same for high-beta. Only traces with significant beta peak were included in the frequency models. [a] Time x DRT estimated coefficient represents time effect given on DRT condition, in addition to the main time effect. A significant positive estimated coefficient indicated that time slope in the on DRT condition was significantly more positive (or less negative) than time slope in the off DRT condition, and vice versa for negative values.
